# Siderophore-mediated zinc acquisition enhances enterobacterial colonization of the inflamed gut

**DOI:** 10.1101/2020.07.20.212498

**Authors:** Hui Zhi, Judith Behnsen, Allegra Aron, Vivekanandan Subramanian, Janet Z. Liu, Romana R. Gerner, Daniel Petras, Keith D. Green, Sarah L. Price, Jose Camacho, Hannah Hillman, Joshua Tjokrosurjo, Nicola P. Montaldo, Evelyn Hoover, Sean Treacy-Abarca, Benjamin A. Gilston, Eric P. Skaar, Walter J. Chazin, Sylvie Garneau-Tsodikova, Matthew B. Lawrenz, Robert D. Perry, Sean-Paul Nuccio, Pieter C. Dorrestein, Manuela Raffatellu

**Affiliations:** Division of Host-Microbe Systems & Therapeutics, Department of Pediatrics, University of California San Diego, La Jolla, CA 92093, USA; Department of Microbiology & Molecular Genetics, University of California Irvine, Irvine, California, USA; Skaggs School of Pharmacy and Pharmaceutical Sciences, University of California San Diego, La Jolla, California, USA; Collaborative Mass Spectrometry Innovation Center, University of California, San Diego, La Jolla, CA 92093, United States of America; University of Kentucky PharmNMR Center, College of Pharmacy, University of Kentucky, Lexington, KY, 40536-0596, USA; Department of Pharmaceutical Sciences, College of Pharmacy, University of Kentucky, Lexington, KY, 40536-0596, USA; Department of Microbiology and Immunology, University of Louisville School of Medicine, Louisville, KY, 40202, USA; Department of Biochemistry and Chemistry, and Center for Structural Biology, Vanderbilt University Medical Center, Nashville, Tennessee, USA; Department of Pathology, Microbiology, and Immunology, Vanderbilt University Medical Center, Nashville, Tennessee, USA; Center for Predictive Medicine for Biodefense and Emerging Infectious Diseases, Department of Microbiology and Immunology, University of Louisville School of Medicine, Louisville, KY, 40202, USA; Department of Microbiology and Immunology, University of Kentucky, Lexington, Kentucky 40536, USA; Center for Microbiome Innovation, University of California San Diego, La Jolla, CA 92093, USA; Chiba University-UC San Diego Center for Mucosal Immunology, Allergy, and Vaccines (CU-UCSD cMAV), La Jolla, CA 92093, USA

## Abstract

Zinc is an essential cofactor for bacterial metabolism, and many *Enterobacteriaceae* express the zinc transporters ZnuABC and ZupT to acquire this metal in the host. Unexpectedly, the probiotic bacterium *Escherichia coli* Nissle 1917 exhibited appreciable growth in zinc-limited media even when these transporters were deleted. By utilizing *in vitro* and *in vivo* studies, as well as native spray metal infusion mass spectrometry and ion identity molecular networking, we discovered that Nissle utilizes yersiniabactin as a zincophore. Indeed, yersiniabactin enables Nissle to scavenge zinc in zinc-limited media, to resist calprotectin-mediated zinc sequestration, and to thrive in the inflamed gut. Moreover, we discovered that yersiniabactin’s affinity for iron or zinc changes in a pH-dependent manner, with higher affinity for zinc as the pH increased. Altogether, we demonstrate that siderophore metal affinity can be influenced by the local environment and reveal a mechanism of zinc acquisition available to many commensal and pathogenic *Enterobacteriaceae*.

## INTRODUCTION

The *Enterobacteriaceae* are a diverse family of bacteria that inhabit the gastrointestinal tract. Members of this group include the enteric pathogen *Salmonella enterica* serovar Typhimurium (*S.* Typhimurium, or STm), as well as *Escherichia coli*, a species that comprises myriad commensals, pathobionts, and pathogens. Both STm and *E. coli* can colonize the intestine of mammals and thrive in inflammatory conditions ^1–6^. During homeostasis, the gut microbiota is primarily composed of obligate anaerobes belonging to the phyla Bacteroidetes and Firmicutes ^7^. In the inflamed gut, however, the oxidative environment suppresses obligate anaerobes and favors the growth of facultative anaerobes, which include pathogenic and commensal *Enterobacteriaceae* ^1,2,4–6,8^.

One mechanism that enables enterobacterial growth in the inflamed gut is the ability to scavenge metal nutrients. Many biological processes including DNA replication, transcription, respiration, and oxidative stress responses require iron, manganese, cobalt, nickel, copper and/or zinc ^9^. Iron is one of the most abundant transition metal ion in living organisms, and serves as an essential cofactor in central metabolism and respiration ^10,11^. The other most abundant is zinc, which is a cofactor for an estimated 5-6% of all proteins ^12^, and whose functions include acting as the catalytic center in enzymes such as metalloproteases, superoxide dismutases, and metallo-β-lactamases. Thus, bacteria must be able to acquire sufficient amounts of both iron and zinc in order to survive and replicate in a given environment.

Bacteria living inside the human host face particular difficulties in obtaining these metal nutrients. During homeostasis, the availability of such metal ions is actively limited by the host and by the resident microbiota. Moreover, nutrient metal availability is further restricted during inflammation in a process termed “nutritional immunity” ^13^, wherein the host secretes antimicrobial proteins that sequester iron, zinc, and manganese from microbes to limit their growth. We have previously shown that the pathogen STm overcomes host nutritional immunity by obtaining iron, zinc and manganese in the inflamed gut ^1,14–16^. In response to iron limitation, STm secretes enterobactin and salmochelin, which are small iron-scavenging molecules called siderophores ^17,18^. In response to zinc limitation, STm expresses the high-affinity zinc transporter ZnuABC ^15,19,20^. STm also expresses the ZupT permease, which transports zinc and other divalent metal ions ^21,22^. Independently, each of these transporters has been shown to contribute to STm virulence in mouse models of infection ^15,19,20,23,24^.

High-affinity zinc acquisition systems enable microbes to overcome zinc sequestration by the host protein calprotectin (CP), a heterodimer of the S100A8 and S100A9 proteins ^25^. CP constitutes up to 40% of neutrophil cytosolic content ^26^, and the expression of its two subunits can be induced in epithelial cells following stimulation with IL-17 and IL-22 ^1,27^. In the inflamed gut, expression of ZnuABC enables STm to overcome CP-mediated zinc sequestration, outcompete the microbiota, and colonize to high levels ^1,15^.

In addition to STm, other *Enterobacteriaceae* can thrive in the inflamed intestine. One such example is the probiotic bacterium *Escherichia coli* Nissle 1917 (*E. coli* Nissle, or EcN), a strain that was first isolated in WWI from the stool of a soldier who did not develop gastroenteritis during a *Shigella* outbreak ^28^. Since then, EcN has proven to be effective in the treatment and prevention of intestinal disorders including chronic constipation, ulcerative colitis, and infantile diarrhea ^29–32^. Despite being used as a probiotic for nearly a century, the mechanisms through which EcN exerts its protective effects are not completely understood.

Our previous work has demonstrated that EcN utilizes multiple iron uptake systems and secretes antimicrobial proteins known as microcins to outcompete and reduce STm colonization in mouse models of gastroenteritis ^3,33^. As a follow-up to these studies, we initially sought to investigate the contribution of high-affinity zinc transporters to gut colonization by EcN. Unexpectedly, we found that an EcN strain lacking ZnuABC and ZupT was still able to grow appreciably in zinc-limited media, leading us to hypothesize that EcN expresses an additional means of acquiring zinc. After a genome search did not yield any promising candidate transporters, we hypothesized that EcN produces and secretes an unknown zincophore. Using our recently developed native spray metabolomics approach ^34^, we subsequently discovered that EcN produces the siderophore yersiniabactin (Ybt), which is capable of binding zinc. Moreover, we demonstrate that EcN utilizes Ybt, in addition to the zinc transporters ZnuABC and ZupT, to effectively acquire zinc *in vitro*, to resist the antimicrobial activity of CP, and to colonize the inflamed gut.

## RESULTS

### *E. coli* Nissle is more resistant to calprotectin-mediated zinc sequestration than *S.* Typhimurium

We have previously shown that multiple iron uptake systems enable EcN to colonize the inflamed gut and to compete with STm ^33^. As zinc is also limited in the inflamed gut, we hypothesized that EcN must also have robust mechanisms for acquiring this metal. We thus compared the growth of EcN to the growth of STm in a rich medium supplemented with CP, a host antimicrobial protein that sequesters zinc and limits its availability to microbes ^15,25^. To this end, we employed CP concentrations (150-250 μg/ml) comparable to those found in the inflamed gut ^15^. EcN and STm grew to the same density after 16 h of culture without the addition of CP, or in the presence of 150 μg/ml CP (**Fig. 1a**). However, we noticed that in media containing CP at 250 μg/ml, EcN grew ∼8 times better than STm (**Fig. 1a**). These results indicated that EcN is more resistant than STm to the antimicrobial activity of CP *in vitro* and prompted us to investigate the underlying mechanism.

**Figure 1.**
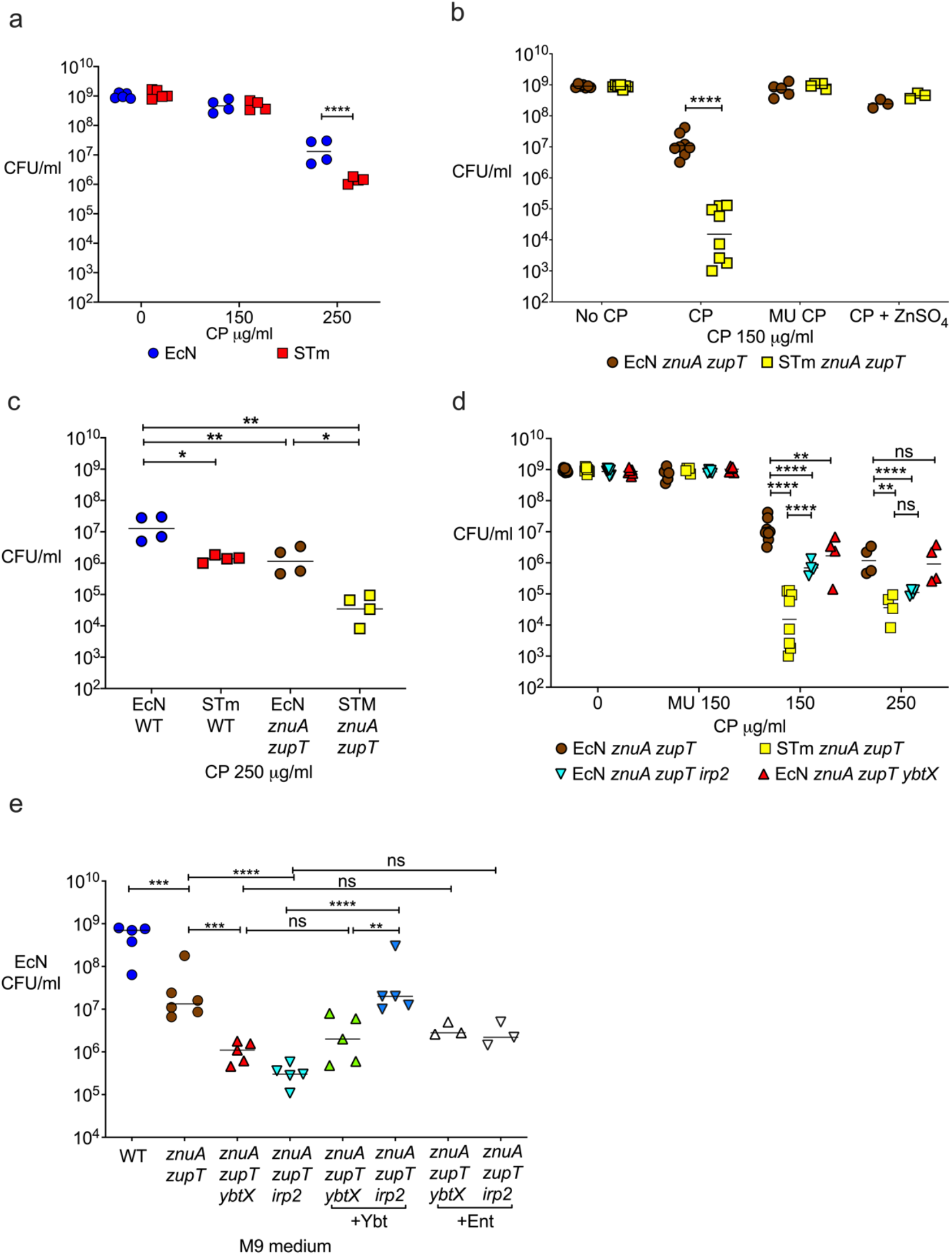
*E. coli* Nissle resistance to calprotectin-mediated zinc limitation *in vitro* is dependent on ZnuABC, ZupT, and a product of the yersiniabactin operon. (**a**) EcN and STm wild-type were grown in modified LB medium without CP, or supplemented with 150 µg/ml or 250 µg/ml CP. (**b**) EcN and STm *znuA zupT* mutants were grown in modified LB medium without CP (No CP), or supplemented with 150 µg/ml CP, 150 µg/ml Site I/II knockout mutant CP (MU CP), or 150 µg/ml CP plus 5 µM ZnSO_4_ (CP + ZnSO_4_). (**c**) EcN and STm wild-type and *znuA zupT* mutants were grown in modified LB medium supplemented with 250 µg/ml CP. (**d**) EcN and STm *znuA zupT* mutants, as well as EcN triple mutants (*znuA zupT irp2*; *znuA zupT ybtX*), were grown in modified LB supplemented with either 150 µg/ml mutant CP (MU 150), or with 150 µg/ml (150) or 250 µg/ml CP (250). (**e**) EcN wild-type and indicated double and triple mutants were grown in M9 medium or in M9 supplemented with either 1 µM yersiniabactin (Ybt) or enterobactin (Ent). (**a-e**) Growth was quantified by enumeration of bacterial CFU on selective media after 16 h static (**a-d**) or 20 h shaking (**e**) incubation. Data are representative of three independent experiments. Bars represent the geometric mean. * *P* value ≤0.05; ** *P* value ≤0.01; *** *P* value ≤0.001, **** *P* value ≤0.0001; ns = not significant.

Both EcN and STm encode two known zinc transport systems: the high affinity zinc transporter ZnuABC and the permease ZupT ^15,19,20,23,24^. Although the function of these two transporters in EcN has not been directly investigated, their disruption significantly diminishes the capacity of the closely-related uropathogenic *E. coli* strain CFT073 to grow in zinc-depleted culture media and to cause urinary tract infection ^35^. To determine whether the difference in CP-resistance between EcN and STm is the result of variations related to ZnuABC and ZupT, we disrupted these transporters in both EcN and STm by deleting the genes *znuA* and *zupT*. As expected, both mutant strains (EcN *znuA zupT* and STm *znuA zupT*) grew slower than their respective parental strains in the presence of CP, but not in the presence a Site I/II knockout mutant CP (MU CP; lacks the ability to bind zinc) ^36,37^, or when ZnSO_4_ was added to the media (**Fig. 1b**). These results indicated that ZnuABC and ZupT have similar functions in both EcN and STm, and mediate evasion of CP-dependent antimicrobial activity.

Puzzlingly, we observed that the EcN *znuA zupT* mutant grew almost 1,000-fold better than the STm *znuA zupT* mutant in the presence of 150 μg/ml CP (**Fig. 1b**). Although higher concentrations of CP (250 μg/ml) reduced the growth of the EcN *znuA zupT* mutant, it was still 100-fold higher than the STm *znuA zupT* mutant (**Fig. 1c**). As addition of ZnSO_4_ rescued the growth of both the EcN and the STm *znuA zupT* mutants (**Supplementary Fig. 1a**), we posited that EcN is able to acquire zinc via an additional mechanism.

### A product of the yersiniabactin operon promotes zinc acquisition by *E. coli* Nissle in zinc-limited media

In iron-limiting conditions, EcN acquires iron by producing the siderophores enterobactin, salmochelin, aerobactin, and Ybt ^33^. Although the importance of siderophores in scavenging iron has been well-demonstrated in biological systems, chemists have known for decades that some siderophores can bind other metals besides iron (reviewed in ^38^). Among the siderophores produced by EcN, Ybt has been shown to also bind copper, gallium, nickel, cobalt, and chromium ^39^. Intriguingly, a product of the Ybt gene cluster has been proposed to contribute to zinc acquisition by the pathogen *Yersinia pestis* ^40,41^; however, its identity and mechanism are unknown, as two prior studies did not provide evidence of direct zinc binding by Ybt ^39,41^. We thus sought to determine whether, in addition to ZnuABC and ZupT, EcN uses a product of the Ybt operon to acquire zinc under zinc-limiting conditions.

To this end, we deleted the *ybt* cluster’s *irp2* gene that encodes the synthetase HMWP2, thus rendering EcN unable to synthesize Ybt ^42–44^. We also deleted the *ybtX* gene, which encodes for an inner membrane permease that was found to promote zinc acquisition by *Y. pestis* ^40,41^. Of note, the first published genome sequence of EcN (GenBank CP007799.1, ^45^) indicated that *irp1 and irp2* were disrupted (frameshifted and insertion sequence, respectively), although a recent sequencing effort of our lab’s EcN strain revealed these genes to be intact (GenBank CP022686.1), which is consistent with a prior study showing that EcN produces Ybt ^46^. Next, we tested the growth of EcN strains lacking these genes, in addition to the *znuA zupT* genes, in metal-limiting conditions (M9 minimal medium). Strains lacking *znuA zupT* and either *irp2* or *ybtX* displayed a severe growth defect in M9 minimal medium, growing 1,000-fold less than EcN wild-type and more than 10-fold less than EcN *znuA zupT* (**Fig. 1e**). Furthermore, growth of all mutants was restored in ZnSO_4_-supplemented M9 minimal medium (**Supplementary Fig. 1b**) and in LB broth without metal limitation (**Supplementary Fig. 1c**), confirming that the observed growth defects were indeed due to zinc deficiency. Taken together, these results suggested that a product of the *ybt* gene cluster contributes to zinc acquisition by the probiotic EcN in zinc-limited media. We therefore hypothesized that the Ybt locus may encode for the production of a zincophore.

### Yersiniabactin is a zincophore

To identify whether the *ybt* gene cluster produces a zincophore, we cultured EcN wild-type and the *irp2* mutant in M9 minimal media and collected culture pellets and supernatants to then run ultra-high performance liquid chromatography tandem mass spectrometry (UHPLC-MS/MS). In addition to running UHPLC-MS/MS metabolomics on these samples, we performed experiments using post-liquid chromatography (LC) pH adjustment to 6.8 and infusion of a zinc acetate solution, followed by mass spectrometry, in order to assess whether any of the metabolites produced were capable of binding zinc^34^. This native spray metabolomics strategy is then combined with ion identity-based molecular networking, a new computational and data visualization strategy that allows for the discovery of mass spectrometry features with the same retention time and predictable mass offsets ^34^; mass spectrometry features with the same retention time and a mass difference resulting from zinc binding can be discovered directly from complex metabolomics samples. Using this native spray metabolomics workflow, a number of zinc-binding small molecules were observed in the wild-type supernatant samples (**Fig. 2a, b**); zinc-bound nodes (each node represents an MS1 feature and its clustered MS/MS spectra) are shown in salmon and are connected to protonated nodes (dark blue) with a blue dashed line (indicating a *m/z* delta = [Zn^2+^-H^+^]^+^) (**Fig. 2a, b**). Furthermore, two peaks were observed in both culture supernatant (**Fig 2c**) and pellets from EcN wild-type that were absent in the *irp2* mutant cultured in M9 minimal media (**Fig. 2c**). Feature-based molecular networking using MZmine 2 in conjunction with Global Natural Products Social (GNPS) Molecular Networking ^47,48^ allowed us to putatively identify these two peaks as Ybt. Ybt is known to tautomerize at C10 (**Fig. 3**) and Ref. ^49^. We confirmed that these peaks were two diastereomers of Ybt by matching the retention time, exact mass, and MS/MS spectra acquired from culture extracts to an authentic Ybt standard (**Fig. 2e, f**). Post-LC pH neutralization and zinc-infusion revealed the zinc-bound Ybt species, indicating that Ybt is indeed capable of binding zinc (**Fig. 2d**). To our surprise, we also found that one of the diastereomers (at retention time = 4.0 min) binds zinc with higher preference than the other (at retention time = 4.3 min) (**Fig. 2d**). Since Ybt was initially discovered as an iron-binding molecule, and thus termed a siderophore, we next sought to determine the preferential conditions for binding iron versus zinc. To assess the competition between iron and zinc binding, we performed direct infusion mass spectrometry competition experiments at multiple pH values. In these experiments, we added equimolar amounts of zinc and iron to Ybt in ammonium acetate buffer adjusted to pH 4, 7, and 10. While Ybt preferentially bound iron at low pH (pH 4), it exhibited a higher preference for zinc at high pH (pH 10) (**Fig. 2g**). Intriguingly, at neutral pH (pH 7), Ybt was observed bound to iron or zinc at roughly equal proportion (**Fig. 2g**).

**Figure 2.**
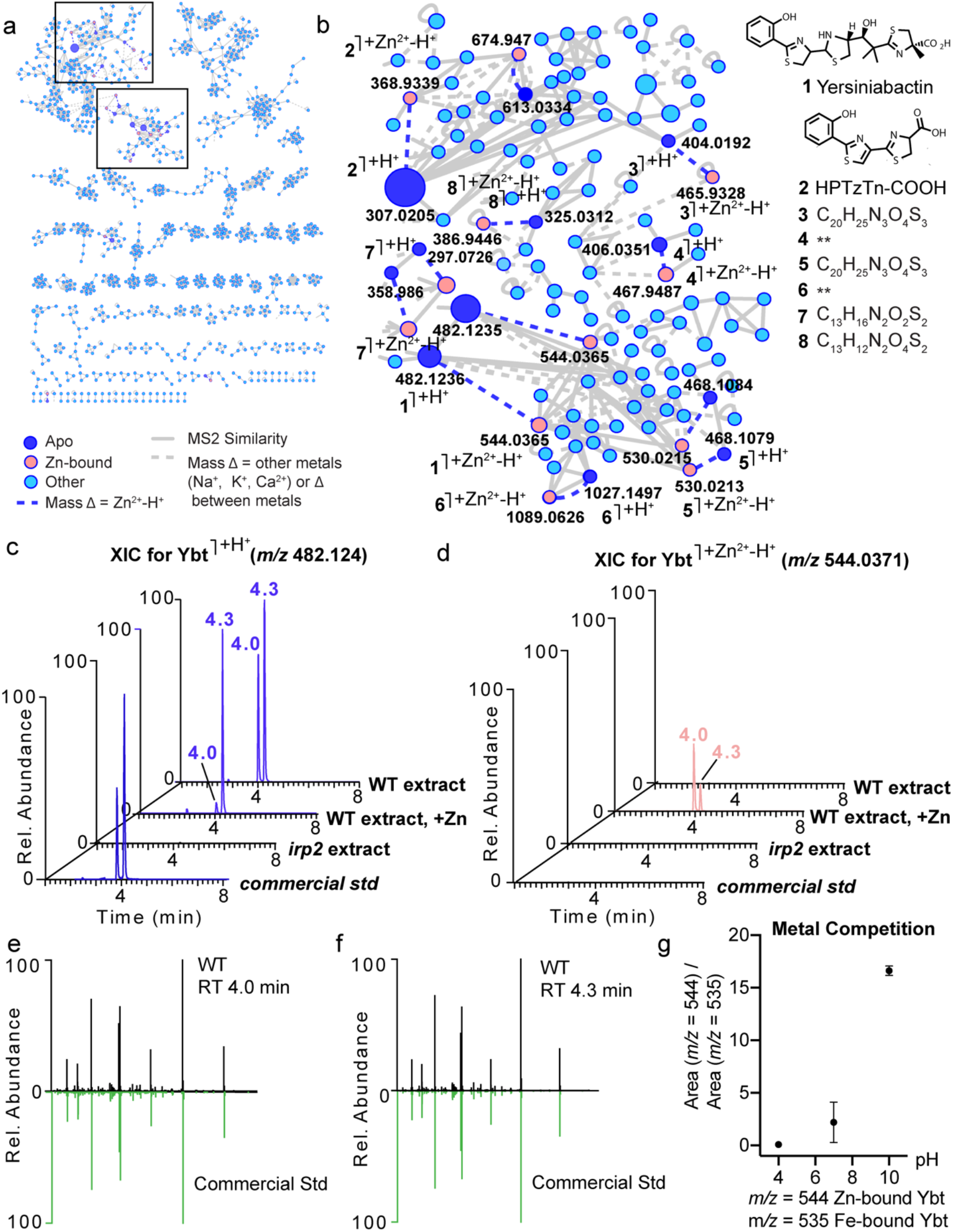
Yersiniabactin is produced by *E. coli* Nissle and directly binds zinc in a pH-dependent manner. (**a, b**) Native spray metal metabolomics was used to identify zinc-binding small molecules present in isolated EcN supernatant extracts. Zinc-binding molecules, including Ybt and various other truncations, are concentrated in the boxed molecular families enlarged in panel B. Zinc-binding small molecules are observed when post-LC infusion of Zn^2+^ and post-pH adjustment are performed. Zinc-bound molecules are shown in salmon, while the corresponding protonated (Apo) form of these molecules is shown in dark blue. Structures and molecular formulas (generated using SIRIUS 4.0 ^78^) are provided. (**c**) Extracted ion chromatogram (XIC) for apo-Ybt ([M+H^+^]^+^ = 482.1236) is observed as two peaks (present at 4.0 and 4.3 minutes) in wild-type EcN (WT extract); tautomerization occurring at C10 results in the racemic mixture (Fig. 3). When post-LC pH adjustment and Zn^2+^-infusion are performed, the majority of the peak at 4.3 remains apo-Ybt (WT extract, +Zn). Neither apo-Ybt peak is present in the EcN *irp2* knockout supernatant (*irp2* extract). Commercial Ybt (*commercial std*) also elutes with a minor peak at 4.0 and a major peak at 4.3 minutes. (**d**) Zn^2+^-bound Ybt ([M+Zn^2+^-H^+^]^+^ = 544.0371) is not observed in the XIC of wild-type samples (WT extract), in *irp2* knockout samples (*irp2* extract), or in the commercial standard (*commercial std*) when the standard LC-MS/MS method is applied; however, when native spray metal metabolomics is applied (WT extract, +Zn; post-LC infusion of Zn^2+^ in conjunction with pH neutralization), Zn^2+^-bound Ybt is observed ([M+Zn^2+^-H^+^]^+^ = 544.0371) as the major species in the first peak (at retention time = 4.01 minutes). (**e, f**) Mirror plots show that peaks present at (**e**) 4.0 min and (**f**) 4.3 min (black; MS/MS of [(M+2H^+^)/2 = 241.5654) both match the MS/MS from commercial Ybt standard (green). (**g**) Metal competition data for direct injection experiments run at pH 4, 7, and 10, in which Ybt was added to buffer in the presence of both iron and zinc. The ratio of extracted peak area of Zn^2+^-bound Ybt ([M+Zn^2+^-H^+^]^+^ = 544.0371) to extracted peak area of Fe^3+^-bound Ybt ([M+Fe^3+^-H^+^]^+^ = 535.0351) is shown at each of the three tested pH values.

**Figure 3.**
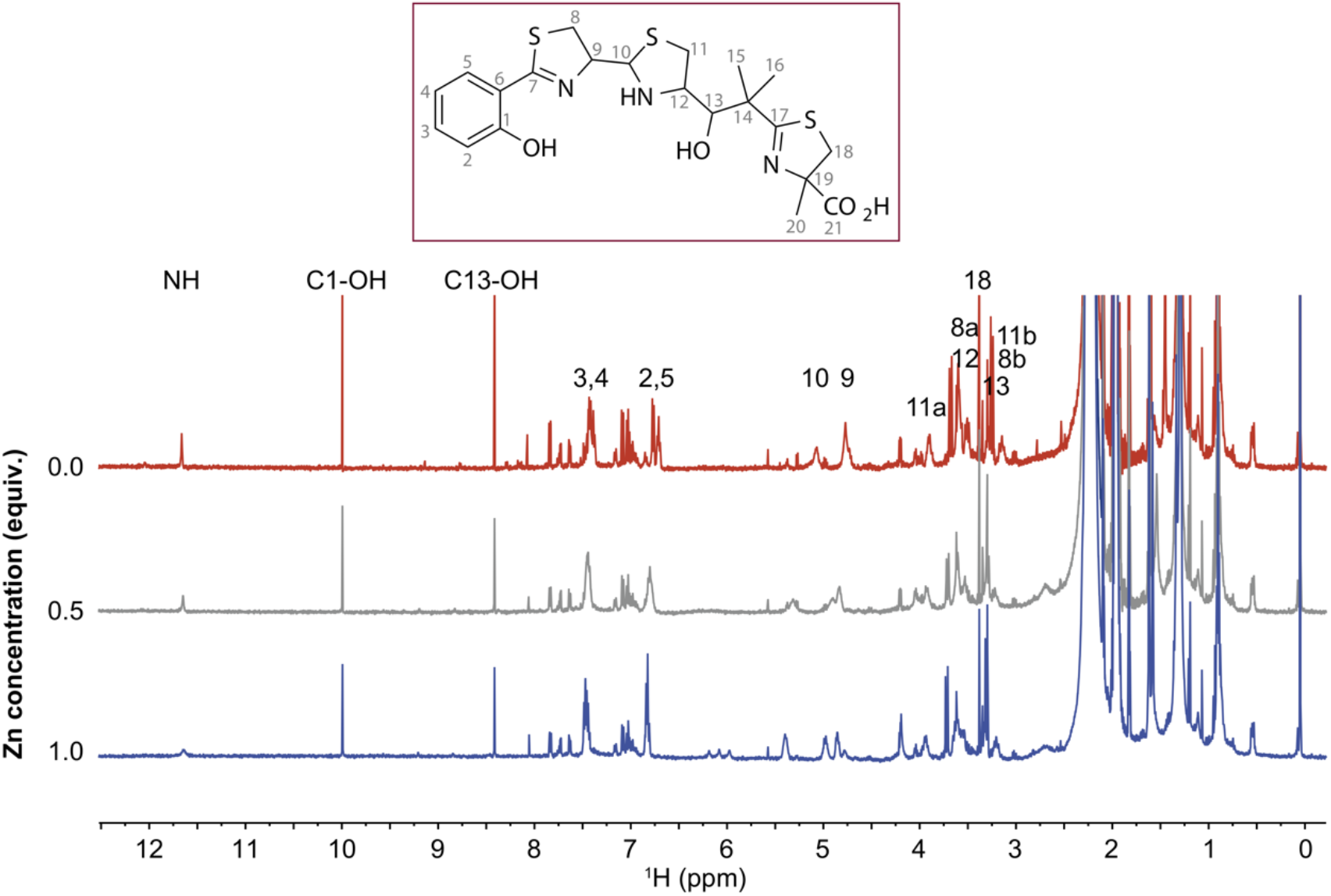
1D ^1^H NMR confirms direct zinc binding to yersiniabactin. 1D ^1^H NMR spectra of Ybt dissolved in CD_3_CN (top trace, 0 equivalents of Zn, red trace) as increasing zinc is titrated into the solution (0.5 equiv., gray trace), (1.0 equiv., blue trace). The loss of intensity of the NH and OH signals arises from the coordination of zinc by the corresponding N and O atoms; the signals that shift correspond to protons whose electronic environment changes due to binding of the Zn^2+^ ion. Only partial binding is observed because Ybt is in rapid equilibrium between two tautomers at C10 in addition to hydrolysis that occurs at C10 ^49^.

To confirm the zinc-binding observed by native electrospray metabolomics, we monitored a zinc-titration into Ybt by 1D ^1^H NMR (**Fig. 3**). Although Ybt is in equilibrium between two tautomers at C10 that have different relative affinities for zinc, only one set of signals is observed in the spectra. This same observation was reported in earlier studies of gallium binding to Ybt ^49^. The addition of zinc modulates a number of signals, including those of the NH proton and the two hydroxyl proton peaks at ∼11.6, 10.0, and 8.4 ppm. The intensity of these well resolved peaks decreases progressively upon addition of 0.5 and 1.0 equivalents of zinc, which is consistent with the nitrogen (N10-12) and oxygen (O1 and O13) heteroatoms chelating the zinc atom (**Fig. 3**), in a manner similar to Ybt binding of iron ^50^ and copper ^51^, and zinc binding by the *Pseudomonas* sp.-derived compound micacocidin A ^52^. Finally, we found that increasing the pH of the solution via addition of 0-5 molar equivalents of NaOD had little effect on the NMR spectrum of zinc-bound Ybt complex, aside from the exchange of labile hydrogens with deuterium (**Supplementary Fig. 2**), showing that zinc is bound in the same manner across a broad range of pH.

Having now discovered that Ybt can bind zinc in a physiologically relevant pH range, we next tested whether the addition of exogenous Ybt could rescue the growth of an EcN strain that is highly susceptible to zinc limitation due to mutations in ZnuABC, ZupT, and Ybt synthesis (*znuA zupT irp2* mutant). Consistent with our hypothesis, supplementation of M9 minimal media with 1 μM purified apo-Ybt (Ybt not bound to iron) rescued the growth of the EcN *znuA zupT irp2* mutant to similar levels as the *znuA zupT* mutant (**Fig. 1e**). Furthermore, the growth of a strain deficient in the putative zinc-transporting inner membrane protein YbtX (*znuA zupT ybtX*) was not significantly rescued by exogenous apo-Ybt (**Fig. 1e**). Addition of the siderophore apo-enterobactin, which is not expected to bind to zinc, did not significantly rescue the growth of either the *znuA zupT irp2* mutant or the *znuA zupT ybtX* mutant (**Fig. 1e**). Taken together, these results demonstrate that Ybt binds to both iron and zinc, that metal binding preference can be influenced by pH, and that Ybt can scavenge zinc for EcN in zinc-limited media. Next, we assessed whether Ybt enables EcN to evade the host response.

### *E. coli* Nissle’s higher resistance to calprotectin is due to yersiniabactin-mediated zinc acquisition

In the host, zinc limitation is largely dependent on the antimicrobial protein CP ^53^. We thus tested whether Ybt-mediated zinc acquisition enhances EcN’s growth in CP-supplemented rich media. Above, we demonstrated that when the ZnuABC and ZupT transporters were deleted (*znuA zupT* mutants), EcN grew better than STm (**Fig. 1b-d**). Moving forward, when either *irp2* or *ybtX* were additionally deleted in EcN, growth of the *znuA zupT irp2* and the *znuA zupT ybtX* mutants were ∼8-fold lower than the parental EcN *znuA zupT* strain in the presence of 150 µg/ml CP (**Fig. 1d**). Although the growth of EcN *znuA zupT* was further diminished in the presence of 250 µg/ml CP, the growth of the EcN *znuA zupT irp2* mutant was again ∼10-fold lower, and now comparable to that of the STm *znuA zupT* mutant (**Fig. 1d**). These results are consistent with Ybt scavenging zinc for EcN when the metal is limited by CP. Because growth of the EcN *znuA zupT ybtX* mutant was similar to the *znuA zupT* mutant in media supplemented with 250 µg/ml CP, it is possible that zinc-bound Ybt can also be internalized via a YbtX-independent mechanism. To confirm that the growth defect of the EcN *znuA zupT irp2* mutant is due to zinc chelation by CP, we supplemented the medium with 150 µg/ml of CP Site I/II knockout mutant (**Fig. 1d**), or with 150 µg/ml CP and 5 µM ZnSO_4_ (**Supplementary Fig. 1a**). In both experiments, all strains grew to the same level. Taken together, these results indicate that Ybt-mediated zinc acquisition enhances EcN resistance to zinc limitation induced by CP and provide a mechanistic explanation for EcN’s heightened resistance to zinc limitation relative to STm.

### Yersiniabactin enhances *E. coli* Nissle colonization of the inflamed gut

After demonstrating that Ybt promotes EcN resistance to CP *in vitro*, we next sought to investigate whether Ybt confers a growth advantage to EcN during inflammatory conditions *in vivo*, where CP is highly expressed ^1,15^ and zinc is limited ^15^. To induce intestinal inflammation, we employed the dextran sodium sulfate (DSS) mouse colitis model (**Fig. 4a**). After 4 days of DSS administration, we orally inoculated the mice with a 1:1 mixture of EcN *znuA zupT* and EcN wild-type, or of EcN *znuA zupT* and one of the EcN triple mutants (*znuA zupT irp2* or *znuA zupT ybtX*).

**Figure 4.**
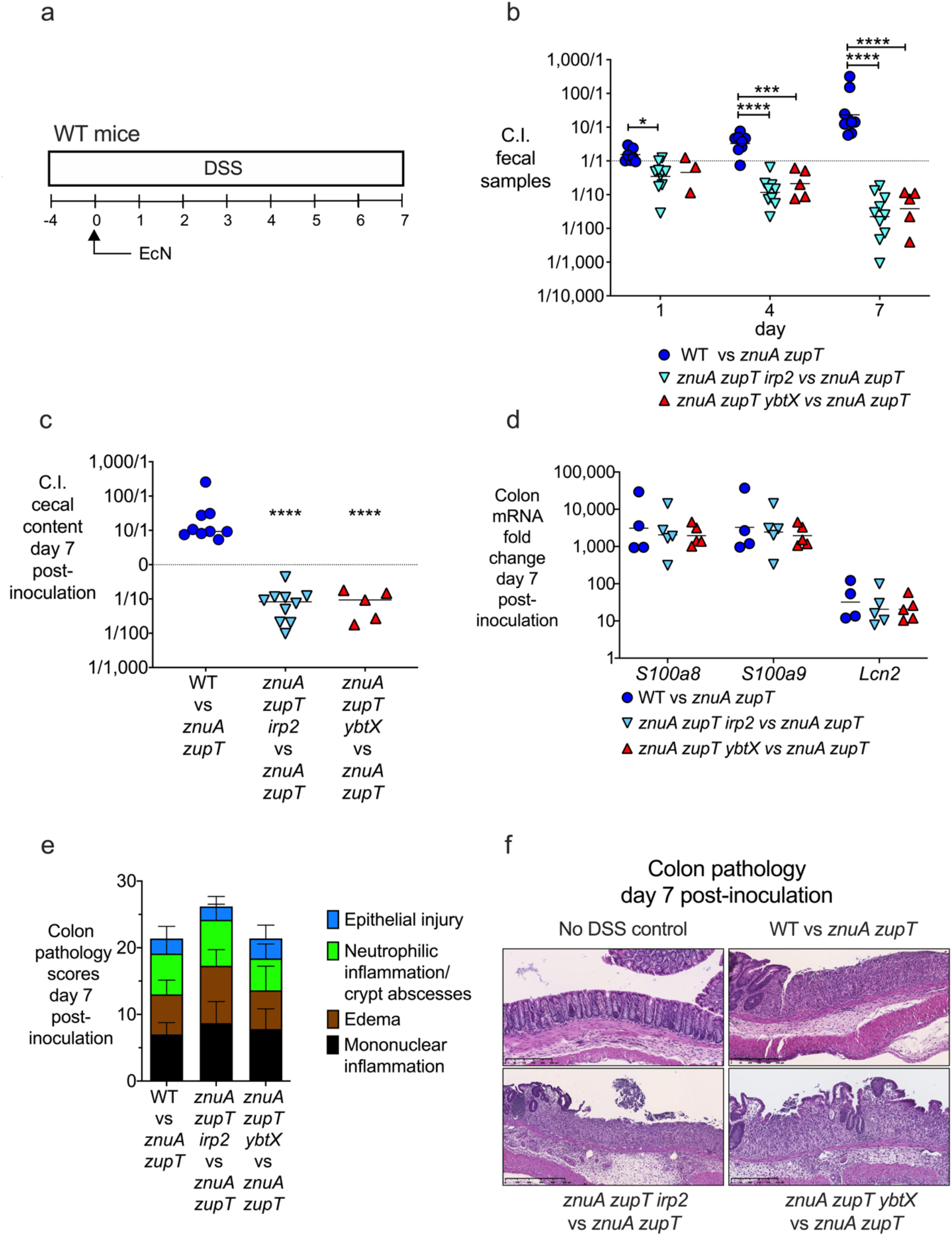
The ability to acquire zinc via yersiniabactin enhances *E. coli* Nissle colonization of the inflamed gut. (a) Experiment timeline for the DSS-induced colitis model and the administration of EcN strains. C57BL/6 mice were given 4% (w/v) DSS in the drinking water for 4 days (day -4 to 0). On day 0, mice were orally gavaged with 1×10^9^ CFU of a 1:1 mixture of EcN strains. (b) Fecal samples were collected on day 1, 4, and 7, and the competitive index (C.I.) was calculated by dividing the output ratio (CFU of either wild type or one of the triple mutants / CFU of the competing *znuA zupT* strain in each group) by the CFU-enumerated input ratio of the strains. (**c**) Cecal content was collected on day 7 and the C.I. of strains in each group was calculated as described in B. (**d**) mRNA expression of *S100a8*, *S100a9* and *Lcn2* was measured in the colon of mice in panel C. (**e**) Colon pathology score of mice in panel C, with sub-scores of each criterion. (**f**) Representative stained sections (H&E, original magnification ×10) of distal colon from healthy or DSS-treated mice administered with different groups of EcN. (**b-d**) Each data point represents a single mouse. Bars represent the geometric mean. (**e**) Bars represent the mean. * *P* value ≤0.05; *** *P* value ≤0.001, **** *P* value ≤0.0001.

EcN wild-type exhibited a significant competitive advantage over the *znuA zupT* mutant beginning at day 1 post-inoculation, which increased to an average of ∼28-fold by day 7 (**Fig. 4b**). These results indicated that ZnuABC and ZupT are needed for optimal colonization of the inflamed gut. By contrast, EcN *znuA zupT* showed a significant competitive advantage over both triple mutants, which increased over time up to ∼20-fold (*znuA zupT ybtX* mutant) and ∼50-fold (*znuA zupT irp2* mutant) (**Fig. 4b and 4c**). In both cases, the increased competitive advantage was due to the decreased colonization level of the triple mutants, as the *znuA zupT* mutant colonized at similar levels (**Supplementary Fig. 3a-c**). Of note, host antimicrobial gene expression levels (*Lcn2*, *S100a8, S100a9*) were similarly upregulated in all DSS-treated mice (**Fig. 4d**), and all DSS-treated mice developed similar levels of colitis, as shown by histopathology evaluation of the distal colon (**Fig. 4e, f**). Collectively, these results indicate that both Ybt production (via Irp2) and Ybt transport (via YbtX) enhance EcN colonization of the inflamed gut. Because Ybt production and acquisition conferred a colonization advantage to the *znuA zupT* mutant, these data support the idea that Ybt can scavenge zinc *in vivo*, in zinc-limited conditions such as those found in the inflamed gut.

### Inflammation and calprotectin are necessary for yersiniabactin to enhance gut colonization by *E. coli* Nissle

Next, we ascertained whether the zinc transport systems of EcN play a significant role in the absence of gut inflammation. As EcN colonization levels decline over time in conventional mice in the absence of inflammation, we used germ-free mice (**Fig. 5a**), in which we previously observed high levels of EcN colonization for extended periods of time ^3^. When we inoculated germ-free mice with a 1:1 mixture of EcN *znuA zupT* and *znuA zupT ybtX* (**Fig. 5a**), we recovered similar amounts of both strains from mouse feces throughout the experiment (**Fig. 5b and Supplementary Fig. 3d**). Whereas *S100a8*, *S100a9*, and *Lcn2* were highly expressed in the ceca of DSS-treated animals colonized with EcN, these genes were only minimally upregulated (less than 10-fold) in germ-free mice colonized with EcN (**Fig. 5c**). The absence of inflammation in EcN*-*colonized germ-free mice was also confirmed by cecal pathology (**Supplemental Fig. 3f,** left panel).

**Figure 5.**
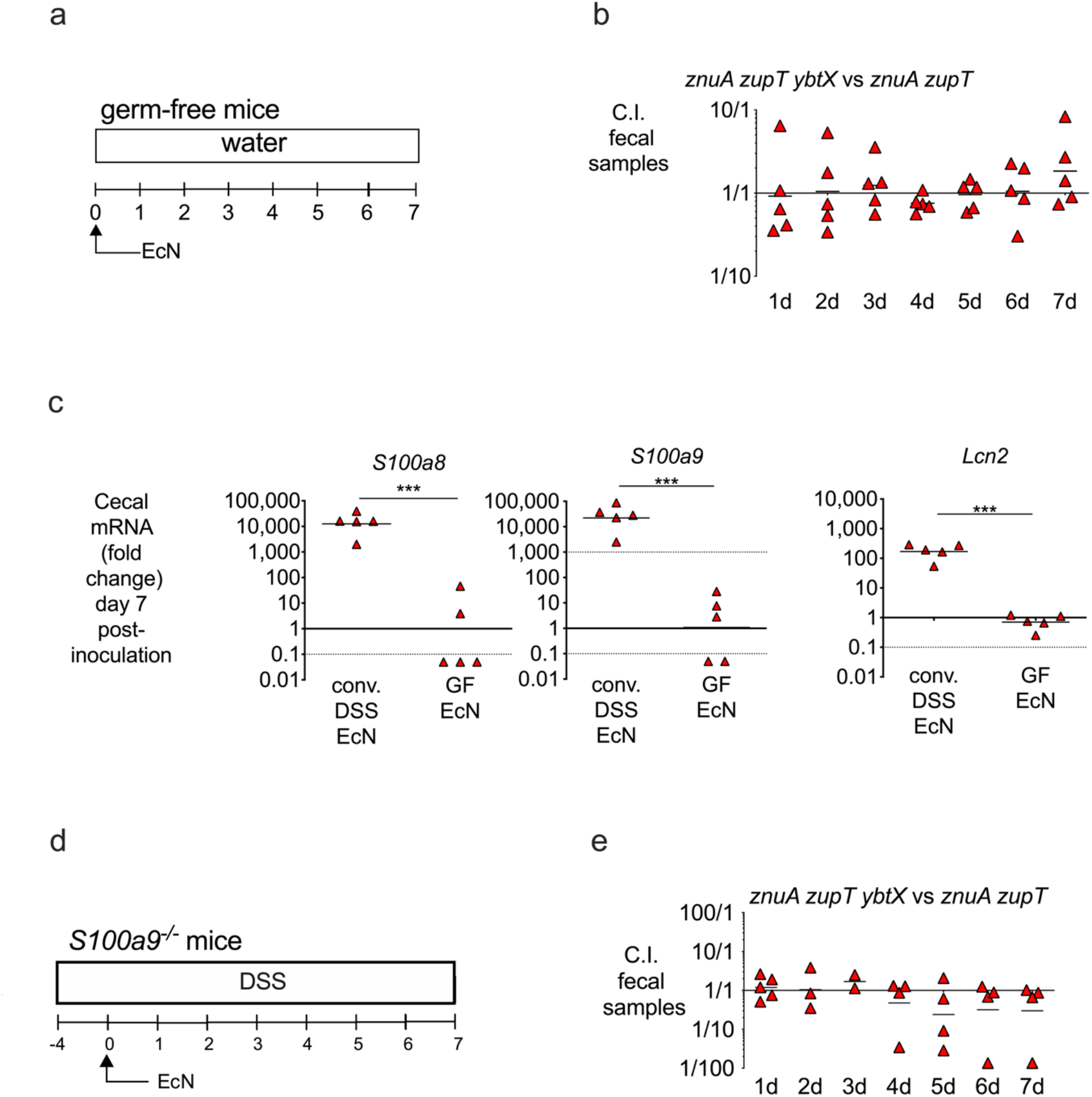
Yersiniabactin-mediated zinc acquisition provides a competitive advantage for *E. coli* Nissle in the presence of inflammation and calprotectin. (**a**) Experiment timeline for panel B. (**b**) Germ-free Swiss Webster mice were colonized with 1×10^9^ CFU of a 1:1 mixture of EcN *znuA zupT* and *znuA zupT ybtX*. Fecal samples were collected daily and the competitive index (C.I.) was calculated by dividing the output ratio (CFU of EcN *znuA zupT ybtX* / CFU of EcN *znuA zupT*) by the CFU-enumerated input ratio of the strains. (**c**) mRNA expression of *S100a8*, *S100a9* and *Lcn2* was measured in the cecum of mice in panel B; conventional DSS-treated mice colonized with EcN were used as a control. (**d**) Experiment timeline for panel E. (**e**) *S100a9*^-/-^ mice were given 4% (w/v) DSS in the drinking water for 4 days (day -4 to 0). On day 0, mice were orally gavaged with 1×10^9^ CFU of a 1:1 mixture of EcN *znuA zupT* and *znuA zupT ybtX*. Fecal samples were collected daily and the C.I. was calculated as described for panel B. Each data point represents a single mouse. Bars represent the geometric mean. *** *P* value ≤0.001.

To further probe whether Ybt provides a means for EcN to evade CP-dependent zinc depletion *in vivo*, we employed *S100a9^-/-^* mice (deficient in CP) treated with DSS, and inoculated them with a 1:1 mixture of EcN *znuA zupT* and *znuA zupT ybtX* (**Fig. 5d**). Although the mice developed intestinal inflammation (**Supplementary Fig. 3f**, right panel), similar amounts of each strain were recovered from these mice lacking CP (**Fig. 5e and Supplementary Fig. 3e)**. Taken together, these results indicate that Ybt confers a colonization advantage to EcN in the inflamed gut, by enabling EcN to evade CP-dependent zinc sequestration.

## Discussion

Commensal and pathogenic *Enterobacteriaceae* exploit host inflammation to achieve high levels of colonization and outcompete obligate anaerobes; these mechanisms include the ability to utilize alternative electron acceptors that become available following the production of reactive oxygen and nitrogen species by activated host cells ^5,6^, as well as new nutrient sources such as lactate ^54^ and acidic sugars ^55^. In addition to taking advantage of new metabolic resources, *Enterobacteriaceae* must also overcome host-mediated mechanisms of nutritional immunity, including metal ion starvation ^13^.

We have previously shown that pathogenic STm and probiotic EcN evade lipocalin-2-mediated iron sequestration in the inflamed gut via the production of stealth siderophores ^16,33^. As we have found that STm also evades CP-mediated zinc sequestration in the inflamed gut ^15^, we sought to investigate whether EcN also evades CP to acquire zinc and thrive in the host. As EcN, akin to STm, expresses ZnuABC and ZupT, we initially hypothesized that these zinc transporters mediate EcN resistance to CP. However, when we found that an EcN *znuA zupT* mutant still grew up to 1,000-fold better than an STm *znuA zupT* mutant in media containing CP (**Fig. 1**), we speculated that EcN must utilize additional mechanisms to acquire zinc. In the work presented herein, we unexpectedly discovered that EcN scavenges zinc with the siderophore Ybt.

Ybt is a phenolate siderophore that was first discovered as being produced by *Yersinia enterocolitica* ^56^. The term siderophore has its origin in the Greek language and means “iron carrier”, as these molecules are widely characterized as being produced by microorganisms in order to acquire iron. However, recent studies have proposed that at least some siderophores may also bind to other metals. For example, the siderophore ferrioxamine was shown to bind manganese ^57,58^, and Ybt was shown to bind copper as a means to evade toxicity ^59^ and to scavenge copper *in vitro* ^60^. Nevertheless, the extent and biological relevance for siderophores binding to other metals remains largely unknown.

Most of the genes involved in Ybt biosynthesis are grouped in a gene cluster ^61,62^. In addition to *Yersinia* species, many *Enterobacteriaceae* also produce Ybt, including both pathogenic and commensal *E. coli* ^46,63–65^. Ybt is well known for scavenging iron *in vivo* ^66^, and plays a critical role in *Y. pestis* virulence ^61^. Moreover, Ybt reduces reactive oxygen species formation in phagocytes by scavenging iron and preventing Haber-Weiss reactions ^67^, as well as contributes to intestinal fibrosis ^63^, indicating that Ybt modulates the host immune response. Incidentally, a product of the *ybt* gene cluster has been proposed to enable zinc acquisition by *Y. pestis* ^40^, although direct binding of Ybt to zinc was not described in two independent studies ^39,41^. Our finding that pH influences binding of Ybt to zinc is likely a key reason for the lack of binding that was observed in these prior publications, as they did not assess changing the pH. Moreover, reinterpretation of the original NMR and UV data in the aforementioned studies does suggest that at least partial zinc coordination can be seen, as the data show slight UV and NMR shifts that are consistent with only a small amount of Ybt being bound to zinc. Nevertheless, a critical question remained as to the identity of the molecule(s) produced by the Ybt gene cluster that contributed to zinc acquisition and, in the context of our study, whether such a molecule could play a role in gut colonization.

Using UHPLC-MS/MS, we identified two diastereomers of Ybt from EcN wild-type supernatant extract that were not present in the *irp2* mutant supernatant; MS/MS spectra of both peaks matched the MS/MS spectrum of commercial Ybt. Ybt is known to isomerize at the C10 position (**Fig. 3**) into a racemic mixture ^49^. Using post-LC pH neutralization and metal infusion in a recently developed workflow termed native metabolomics, we found that one isomer (retention time = 4.0 min) preferentially binds zinc (**Fig. 2**). The different affinity of siderophore diastereomers for a metal is not unprecedented. Pyochelin, a siderophore with a similar thiazoline core as Ybt and produced by *Burkholderia cepacia* and several *Pseudomonas* strains, also exists as two diastereomers, only one of which binds iron ^68^. Moreover, although pyochelin was shown to bind both iron and zinc *in vitro* ^69^, to our knowledge, the biological relevance of pyochelin-mediated zinc scavenging has not been investigated. Similarly, only one of the Ybt isomers was shown to bind gallium when the compound’s structure was initially characterized ^49^. We used 1D ^1^H NMR spectroscopy to confirm this observed zinc binding by Ybt (**Fig. 3**). Specifically, we observed the loss of signal corresponding to the two OH groups and the NH proton in Ybt, which indicates the corresponding heteroatoms that chelate the Zn^2+^ ion. Although the other zinc-coordinating atoms were not directly characterized, we hypothesize that Ybt binds zinc in the same manner as it binds iron and as micacocidin A binds Zn^2+^.

Because Ybt is known to bind iron, we performed a competition assay with equimolar amounts of zinc and iron. We observed that the metal-binding preference of Ybt is pH-dependent – Ybt preferentially binds to zinc in basic conditions (pH = 10), to iron in acidic conditions (pH = 4), and exhibits similar preference for both at pH 7 (**Fig. 2e-h**). In contrast to Ybt, the binding capacity of pyochelin to different metals is pH independent ^69^. We speculate that the pH-dependent metal selectivity of Ybt may confer different functions to it under various physiological conditions (*e.g.*, inflammation or homeostasis), or different colonization niches. For instance, in healthy human subjects, the pH in the small intestine gradually increases from pH 6 in the duodenum to about pH 7.4 in the terminal ileum, then drops to pH 5.7 in the caecum, but again gradually increases, reaching pH 6.7 in the rectum ^70^. Upon inflammation, the pH in most sections of the gastrointestinal tract further decreases, but the colon still possesses a higher pH than the small intestine and cecum. Because dietary iron is mainly absorbed in the small intestine, the ability of Ybt to bind iron at lower pH may enable EcN and other Ybt-producing bacteria to compete with the host for iron in the small intestine. On the other hand, the zinc-binding ability of Ybt may enhance colonization of Ybt-producing bacteria in the colon, where the pH is higher. Intriguingly, in patients with active inflammatory bowel disease, the pH in many sections of the intestine increases; for example, the terminal ileum has been observed to reach up to pH 9.2 ^71^.

During colitis, neutrophils are recruited to sites of inflammation and secrete high levels of CP to sequester zinc from invading pathogens ^15,25,72^. Our observation that Ybt renders EcN more resistant than STm to zinc sequestration by CP (**Fig. 1**) *in vitro* prompted us to investigate the function of Ybt during EcN colonization of the inflamed gut. We found that EcN mutants lacking either Ybt or the putative inner membrane receptor YbtX, in addition to lacking ZnuABC and ZupT, showed more severe colonization defects than the *znuA zupT* mutant in mice with DSS-induced colitis (**Fig. 4**). As four other iron transport systems (including the stealth siderophores salmochelin and aerobactin, as well as heme uptake) are still present in these strains, it is unlikely that the *in vivo* phenotype of the mutants is due to an inability to overcome iron starvation.

Together with the observations that Ybt contributes to optimal growth of EcN in zinc-limited conditions *in vitro* (**Fig. 1**), and that Ybt directly binds zinc at the pHs found in the intestine (**Fig. 2**), the colonization defect of EcN *znuA zupT irp2* and of EcN *znuA zupT ybtX* in DSS-treated mice is consistent with the strains’ limited ability to acquire zinc. Moreover, the colonization advantage provided by Ybt is highly dependent on the state of inflammation and presence of CP, as EcN *znuA zupT* and EcN *znuA zupT ybtX* colonized to similar levels in *S100a9^-/-^* mice as well as in germ-free mice (which lack inflammation and only express low levels of CP) (**Fig. 5**). These results are in agreement with the *in vitro* results showing that Ybt and the putative receptor YbtX enable EcN to acquire zinc in media supplemented with CP (**Fig. 1**).

Altogether, our work demonstrates that Ybt directly binds to zinc in a pH-dependent manner, and that EcN can use Ybt in physiologic, zinc-limiting conditions and in the inflamed gut to evade zinc sequestration by CP. Broadly, our study proposes that the role of Ybt and other siderophores may be more complex than previously thought, and may involve scavenging zinc in the host. Because many commensal and pathogenic *Enterobacteriaceae* (including *Yersinia* spp., *E. coli,* and *Klebsiella pneumoniae*) produce Ybt, this important mechanism of zinc acquisition in the gut may also play a role in other host tissues where pathogens must scavenge zinc.

## MATERIALS AND METHODS

### Bacterial strains, plasmids, and growth conditions

Bacterial strains and plasmids are listed in Supplemental Table 1. Cultures of STm and *E. coli* were routinely incubated either aerobically at 37 °C in Lysogeny broth (LB; per liter: 10 g tryptone, 5 g yeast extract, 10 g NaCl) or on LB agar plates (1.5% Difco agar) overnight. Antibiotics and other chemicals were added at the following concentrations (mg/L) as needed: carbenicillin (Carb), 100; chloramphenicol (Cm), 30; kanamycin (Km), 50 or 100; nalidixic acid (Nal), 50; 5-bromo-4-choloro-3-indoyl-ß-D-galactopyranoside (Xgal), 40. For counterselection of pJK611, 10 % (w/v) sucrose was added to media.

### *E. coli* Nissle 1917 mutant generation

Mutants in EcN and STm were constructed using either the lambda Red recombinase system or allelic exchange deletion. To generate mutants with the lambda Red recombinase system ^73^, primers (Supplemental Table 2) homologous to sequences flanking the 5′ and 3′ ends of the target regions were designed and were used to replace the selected genes with a chloramphenicol (derived from pKD3), a kanamycin (derived from pKD4), or a tetracycline resistance cassette (Supplemental Table 2). Strain names for the mutants are listed in Supplemental Table 1. To confirm integration of the resistance cassette and deletion of the target, mutant strains and wild-type controls were each assayed utilizing PCR, and sequencing primers (Supplemental Table 2) that flank the target sequence were used in conjunction with a common test primer to test for both new junction fragments.

### Bacterial Growth in LB, modified LB supplemented with calprotectin, and M9 minimal medium

STm and EcN strains were tested for their ability to grow in nutrient-rich conditions (LB), nutrient-limited conditions (M9 minimal medium per liter; 6.8 g Na_2_HPO_4_, 3 g KH_2_PO_4_, 0.5 g NaCl, 1 g NH_4_Cl, 0.1 mM CaCl_2_, 1 mM MgSO_4_, 0.2 % glucose), and in modified LB supplemented with calprotectin (CP). Bacteria were inoculated into M9 minimal medium from an LB agar plate, then shaken overnight at 37°C. Absorbance (λ = 600 nm) of the overnight cultures was determined by spectrophotometry, 10^9^ colony-forming units (CFU) were harvested by centrifugation, washed with M9 medium twice, then serially diluted in M9. For growth assays, 5 µl of the 10^7^ CFU/ml dilution were used to inoculate 95 μl of LB or M9. Growth was assessed by CFU enumeration on agar plates after incubating cultures for 24 hours at 37 °C with shaking. Growth was also tested in M9 minimal medium supplemented with 5 μM ZnSO_4_, 1 μM apo-yersiniabactin (EMC Microcollections), or 1 μM apo-enterobactin (kindly provided by Dr. Elizabeth Nolan, MIT). For growth in modified LB supplemented with CP, 10 µl of 10^5^ CFU/ml was used to inoculate 90 μl of LB supplemented with CP buffer (20 mM Tris pH 7.5, 100 mM β-mercaptoethanol, 3 mM CaCl_2_) and 150 or 250 μg/ml wild-type CP, or 150 μg/ml Site I/II mutant CP (MU CP), respectively (10:28:62 ratio of inoculum to LB media to CP buffer). CP was produced as described previously ^37^. Growth was assessed by enumerating CFU on agar plates after incubating cultures statically for 16 hours at 37 °C.

### Yersiniabactin standard sample preparation for MS

Yersiniabactin (acquired from EMC Microcollections, https://www.microcollections.de/) stock solutions were prepared by resuspension of the compound in ethanol to a concentration of 1 mM. A 20 µM solution was prepared for mass spectrometry (MS) analysis. Solutions for analysis were prepared in 50 % methanol/50 % water. Additional solutions were prepared in water + 0.1 % formic acid (pH 2.8), in water, or in 10 mM ammonium bicarbonate (pH 7.7).

### *E. coli* Nissle sample preparation for MS

Supernatants from wild-type and *irp2* knockout *E. coli* Nissle cultures were extracted onto pre-washed SPE cartridges. SPE cartridges were activated 3x with MeOH (3x 3 ml), then were washed 2x with water + 0.1% formic acid (3x 3 ml). Sample was loaded dropwise (steady single dripping) onto SPE cartridges, then cartridges were washed with water + 0.1% formic acid (3x 3 ml). Sample was eluted into 1.9 ml MeOH, then this was concentrated by speed evaporation at room temperature. Samples were weighed and reconstituted with 80% MeOH/20% water + 0.1% FA to a final concentration of 1mg/ml. 5 µL of sample were injected per run.

### Direct Inject-MS data acquisition

For MS analysis, 5 µl were injected through a Vanquish UHPLC system into a Q Exactive Orbitrap mass spectrometer (Thermo Fisher Scientific, Bremen, Germany). A flow rate between 0.2 ml/min and 0.4 ml/min was used for experiments. Data acquisition was performed in MS1 in positive mode. Electrospray ionization (ESI) parameters were set to 52 L/min sheath gas flow, 14 L/min auxiliary gas flow, 0 L/min sweep gas flow, and 400 °C auxiliary gas temperature. The spray voltage was set to 3.5 kV and the inlet capillary to 320 °C. 50 V S-lens level was applied. MS scan range was set to 150-1500 m/z with a resolution at m/z 200 (R_m/z 200_) of 35,000 with one micro-scan. The maximum ion injection time was set to 100 ms with an automated gain control (AGC) target of 1.0E6.

### UHPLC-MS/MS data acquisition

For LC-MS/MS analysis, 5 µl were injected into a Vanquish UHPLC system coupled to a Q Exactive Orbitrap mass spectrometer (Thermo Fisher Scientific, Bremen, Germany). For the chromatographic separation, a C18 porous core column (Kinetex C18, 50 x 2 mm, 1.8 um particle size, 100 Angstrom pore size, Phenomenex, Torrance, USA) was used. For gradient elution, a high-pressure binary gradient system was used. The mobile phase consisted of solvent A (H_2_O + 0.1 % FA) and solvent B (acetonitrile + 0.1 % FA). The flow rate was set to 0.5 ml/min. After injection, the samples were eluted with the following linear gradient: 0-0.5 min 5 % B, 0.5-5 min 5-99 % B, followed by a 2 min washout phase at 99% B and a 3 min re-equilibration phase at 5% B. Data-dependent acquisition (DDA) of MS/MS spectra was performed in positive mode. ESI parameters were set to 52 L/min sheath gas flow, 14 L/min auxiliary gas flow, 0 L/min sweep gas flow, and 400 °C auxiliary gas temperature. The spray voltage was set to 3.5 kV and the inlet capillary to 320 °C. 50 V S-lens level was applied. MS scan range was set to 150-1500 m/z with a resolution at m/z 200 (R_m/z 200_) of 35,000 with one micro-scan. The maximum ion injection time was set to 100 ms with an AGC target of 1.0E6. Up to 5 MS/MS spectra per MS1 survey scan were recorded in DDA mode with R_m/z 200_ of 17,500 with one micro-scan. The maximum ion injection time for MS/MS scans was set to 100 ms with an AGC target of 3.0E5 ions and minimum 5% C-trap filling. The MS/MS precursor isolation window was set to m/z 1. Normalized collision energy was set to a stepwise increase from 20 to 30 to 40% with z = 1 as default charge state. MS/MS scans were triggered at the apex of chromatographic peaks within 2 to 15 s from their first occurrence. Dynamic precursor exclusion was set to 5 s. Ions with unassigned charge states were excluded from MS/MS acquisition as well as isotope peaks.

### Post LC-MS/MS pH neutralization and metal addition for native spray mass spectrometry

A stock solution of 160 mM Zn(CH_3_CO_2_)_2_ was prepared, then diluted to a final concentration of 3.2 mM. A stock solution of ammonium hydroxide at 1 M was also prepared. Sample was run through a C18 column at a flow rate of 0.5 ml/min. Before electrospray, a neutralizing solution of 1 M ammonium hydroxide was added at a flow rate of 5 μl/min, then the solution of 3.2 mM zinc acetate was added at a flow rate of 5 μl/min. Post-LC pH was verified by collecting the flow through and spotting on pH paper (Sigma).

### Ion identity molecular networking of wild-type *E. coli* Nissle supernatant extracts with post-LC zinc infusion

MS was run as described in the LC-MS/MS data acquisition section. MS/MS spectra were converted to .mzML files using MSconvert (ProteoWizard)^74^. All raw and processed data is publicly available at ftp://massive.ucsd.edu/MSV000083387/. MS1 feature extraction and MS/MS pairing was performed with MZMine 2.37corr17.7_kai_merge2 ^34,75,76^. An intensity threshold of 1E6 for MS1 spectra and of 1E3 for MS/MS spectra was used. MS1 chromatogram building was performed within a 10 ppm mass window and a minimum peak intensity of 3E5 was set. Extracted Ion Chromatograms (XICs) were deconvoluted using the local minimum search algorithm with a chromatographic threshold of 0.01%, a search minimum in RT range of 0.1 min, and a median m/z center calculation with m/z range for MS2 pairing of 0.01 and RT range for MS2 scan pairing of 0.2. After chromatographic deconvolution, MS1 features linked to MS/MS spectra within 0.01 m/z mass and 0.2 min retention time windows. Isotope peaks were grouped and features from different samples were aligned with 10 ppm mass tolerance and 0.1 min retention time tolerance. MS1 peak lists were joined using an m/z tolerance of 10 ppm and retention time tolerance of 0.1 min; alignment was performed by placing a weight of 75 on m/z and 25 on retention time. Gap filling was performed using an intensity tolerance of 10%, an m/z tolerance of 10 ppm, and a retention tolerance of 0.1. Correlation of co-eluting features was performed with the metaCorrelate module; retention time tolerance of 0.1, minimum height of 1E5, noise level of 1E4 were used. A correlation of 85 was set as the cutoff for the min feature shape corr. The following adducts were searched: [M + H^+^]^+^, [M + Na^+^]^+^, [M + K^+^]^+^, [M + Ca^2+^]^2+^, [M + Zn^2+^-H^+^]^+^, and [M-H2O], with an m/z tolerance of 10 ppm, a maximum charge of 2, and maximum molecules/cluster of 2. Peak areas and feature correlation pairs were exported as .csv files and the corresponding consensus MS/MS spectra were exported as an .mgf file. For spectral networking and spectrum library matching, the .mgf file was uploaded to the feature-based molecular networking workflow on GNPS (gnps.ucsd.edu) ^47,48,77^. For spectrum library matching and spectral networking, the minimum cosine score to define spectral similarity was set to 0.7. The Precursor and Fragment Ion Mass Tolerances were set to 0.01 Da and Minimum Matched Fragment Ions to 4, Minimum Cluster Size to 1 (MS Cluster off). When Analog Search was performed, the maximum mass difference was set to 100 Da. The GNPS job for the siderophore mix can be accessed: https://gnps.ucsd.edu/ProteoSAFe/status.jsp?task=525fd9b6a9f24455a589f2371b1d9540. All .csv and .mgf files in addition to MZmine 2 project can be accessed at ftp://massive.ucsd.edu/MSV000083387. Mgf files were exported for SIRIUS in MZmine2, then molecular formulas were determined using SIRIUS 4.0.1 (build 9) ^78^ and molecular formulas can be accessed: http://gnps.ucsd.edu/ProteoSAFe/status.jsp?task=e2bd16458ec34f3f9f99982dedc7d158.

### Metal competition MS experiments

Commercial yersiniabactin (dissolved to 1 mM in ethanol) was added to 10 mM ammonium acetate buffer at a defined pH (as determined by pH meter, Denver Instrument UltraBasic) for a final concentration of 10 µM. Acetic acid was added to the buffer to lower the pH to 4, and ammonium hydroxide was added to the buffer to raise the pH to 10. Solutions of zinc acetate and iron chloride were prepared to a final concentration of 10 mM in water; from this solution, both iron and zinc were added to a final solution of 100 µM.

### Data sharing

All mass spectrometry .raw and centroid .mzXML or .mzML files, in addition to MZmine 2 outputs and project file, are publicly available in the mass spectrometry interactive virtual environment (MassIVE) under massive.ucsd.edu with project identifier <u>MSV000083387</u> (*E. coli* Nissle siderophores); raw spectra of yersiniabactin commercial standards are available under <u>MSV000084237</u> (Siderophore Standard Mixture with metal additions). Ion Identity Molecular Networks can be accessed through gnps.ucsd.edu under direct links: https://gnps.ucsd.edu/ProteoSAFe/status.jsp?task=525fd9b6a9f24455a589f2371b1d9540 and http://gnps.ucsd.edu/ProteoSAFe/status.jsp?task=e2bd16458ec34f3f9f99982dedc7d158.

### NMR experiments and signal assignments

All NMR experiments were performed on a Varian 500 MHz spectrometer equipped with a ^1^H channel cold-probe. The yersiniabactin (Ybt, 0.25 mg) sample was dissolved in deuterated acetonitrile (CD_3_CN, 300 µl). The NMR spectra were acquired at 298 K in 3-mm NMR tubes and raw data were processed using Bruker Topspin version 4.0.7. The ^1^H peaks resonance assignments were made using a combination of 2D COSY (**Supplementary Fig. 2b**), 2D ROESY with a mixing time of 300 ms (**Supplementary Fig. 2c**), and natural abundance ^1^H-^13^C HSQC (not shown) spectra. The assignments were made with the assistance of NMRFAM-SPARKY. The 1D experiments were acquired with a relaxation delay of 1 sec, 90-degree ^1^H pulses of about 9.0 µs, and a spectral width of 8000 Hz. 2D ROESY spectra were acquired with a spinlock of 200-300 ms, using 128 transients per FID, and 128 points in the indirect dimension. The natural abundance ^13^C-^1^H HSQC spectrum was acquired using phase-sensitive sequence using TPPI gradient selection, optimized for a heteronuclear coupling constant of 145 Hz, shaped pulses for all 180-degree pulses on the ^1^H channel, H decoupling during acquisition, sensitivity enhance acquisition, and gradient back-INEPT.

### Zn binding and base titration for NMR studies

1D ^1^H NMR titration experiments were carried out to investigate the binding of Zn^2+^ (added as ZnCl_2_) to Ybt. Ybt (0.25 mg) was dissolved in deuterated acetonitrile (CD_3_CN, 300 μl) then a baseline spectrum was taken. A spectrum was recorded between each addition of ZnCl_2_. 0.5 equiv. of ZnCl_2_ dissolved in CD_3_CN was added to the dissolved Ybt, followed by a second 0.5 equiv. (1 equiv. total), another 1 equiv. (2 equiv. total), and finally another 3 equiv. (5 equiv. total). Once the spectrum with 5 equiv. of ZnCl_2_ was recorded, an NaOD (in D_2_O) titration was performed to determine the effect of increasing pH on the binding of Zn^2+^. Sequential additions of 0.5 equiv., a second 0.5 equiv., 1 equiv., and 3 equiv. were made and 1D ^1^H NMR spectra were recorded for each titration point.

### Mouse Experiments

Germ-free Swiss Webster mice as well as specific pathogen-free C57BL/6 wild-type mice and *S100a9*^-/-^ mice were used in our study, in accordance with protocols and guidelines approved by the Institutional Animal Care and Use Committee of the University of California, Irvine and the University of California, San Diego. C57BL/6 mice were purchased from Jackson Lab, whereas *S100a9*^-/-^ mice ^79^ were bred in-house. Germ-free Swiss Webster mice were purchased from Taconic Farms and then bred in-house in germ-free isolators (Park Bio). For experiments, germ-free mice were transferred to sterile housing inside a biosafety cabinet, then colonized with the respective bacterial strains. For chemical colitis experiments using dextran sodium sulfate (DSS), mice were administered 4% (w/v) DSS (MP Biomedicals) in the drinking water beginning 4 days prior to administering bacteria, then provided a fresh 4% DSS solution one day prior. On the day of inoculation, mice were switched to 2% (w/v) DSS in the drinking water and orally gavaged with 1×10^9^ CFU of a mixture of strains at a 1:1 ratio, as indicated. A fresh 2% DSS solution was provided on day 4 post-inoculation. At day 7 post-inoculation, mice were humanely euthanized. Fecal content was collected on days 1, 4, and 7, and CFU were enumerated by plating on appropriate selective agar media. In all mixed inoculation experiments, the competitive index of the EcN strains used in each group were calculated. Groups of 5-10 male and female mice were used for each experiment.

### Quantitative real-time PCR

Total RNA was extracted from cecal and colon tissues of wild-type mice, or cecal tissues of germ-free mice, with TRI Reagent (Sigma-Aldrich), followed by processing with an RNeasy Mini Plus kit (Qiagen). For analyzing gene expression by quantitative real-time PCR, cDNA from each RNA sample was prepared with the SuperScript IV VILO Master Mix with ezDNase kit (ThermoFisher). Real-time qPCR was performed with PowerUp SYBR Green Master Mix (ThermoFisher) and a QuantStudio 5 (ThermoFisher). Data were analyzed using the comparative 2^-ΔΔCt^ method. Target gene expression in each tissue sample was normalized to the respective levels of *Actb* mRNA (β-actin), and compared to uninfected samples.

### Histopathology

Distal colonic tissues from wild-type mice, proximal colonic tissues from *S100a9^-/-^* mice, and cecal tissues from germ-free mice were fixed in 10% buffered formalin, then processed according to standard procedures for paraffin embedding. 5 µm sections were stained with hematoxylin and eosin, then slides were scanned on a NanoZoomer Slide scanner (Hamamatsu) and scored in a blinded fashion as previously described ^3^, with minor modifications. Briefly, each of four histological criteria (mononuclear infiltration, edema, epithelial injury, and neutrophilic inflammation/crypt abscesses) was determined as absent (0), mild (1), moderate (2), or severe (3). Furthermore, each parameter was assigned an extent factor reflecting its overall involvement ranging from 1 (<10%), 2 (10-25%), 3 (25-50%), and 4 (>50%). Scores represent the sum of the above scores in colon or cecum sections.

### Statistical Analysis

CFU data were transformed to Log_10_ and passed a normal distribution test before running statistical analyses. CFU from *in vitro* growth experiments and mouse experiments were compared by one-way ANOVA followed by Tukey’s multiple comparisons test. An adjusted *P* value equal to or below 0.05 was considered statistically significant; * indicates a *P* value ≤0.05, ** *P* value ≤0.01, *** *P* value ≤0.001, **** *P* value ≤0.0001. For qPCR data analysis, a multiple *t*-test was performed on gene expression levels between *znuA zupT/znuA zupT ybtX* treated wild-type mice and germ-free mice; * indicates a *P* value ≤0.05, ** *P* value ≤0.01.

## Acknowledgements

Work in MR lab is supported by Public Health Service Grants AI126277, AI114625, AI145325, by the Chiba University-UCSD Center for Mucosal Immunology, Allergy, and Vaccines, and by the UCSD Department of Pediatrics. M.R. also holds an Investigator in the Pathogenesis of Infectious Disease Award from the Burroughs Wellcome Fund. RRG was partly supported by a fellowship from the Max Kade Foundation and by a fellowship from the Crohns and Colitis Foundation. Work in the EPS and WJC laboratories is supported by Public Health Service Grant AI101171. MBL was supported by NIH grants P20GM125504, R21AI119557, and in part by a grant from the Jewish Heritage Fund for Excellence Research Enhancement Grant Program at the University of Louisville School of Medicine. SLP was supported by T32AI132146 and F31AI147404. PCD acknowledges the support by NIH for this work under 5U01AI124316, P41GM103484, GMS10RR029121, and Gordon and Betty Moore Foundation for the development of the computational infrastructure to study symbiotic interactions. The authors would like to thank Joe Burlison in the Medicinal Chemistry Lab at the James Graham Brown Cancer Center at the University of Louisville for help with Ybt purification and the UK PharmNMR Center in the College of Pharmacy at the University of Kentucky for NMR support.

## Author contributions

JB and MR conceived the overall study. JB, JZL, HZ, S-PN, MR designed the *in vitro* growth assays and the *in vivo* experiments and analyzed the data. JB, JZL, HZ, RRG, JC, HH, JT, NPM, EH, and ST-A performed the *in vitro* growth assays and the *in vivo* experiments. AA, DP, and PCD designed the MS experiments and analyzed the data. AA and DP ran the MS experiments. VS performed the NMR experiments and analyzed the data. KDG and SLP performed Ybt extractions for NMR experiments and analyzed the data. SG-T and MBL designed the NMR experiments and analyzed the NMR data. BAG, EPS, and WJC designed *in vitro* growth assays and provided key reagents. RRG analyzed the histopathology. S-PN performed comparative genomic analysis. WJC, AA, PCD, and RDP analyzed and discussed the NMR data. JB, JZL, HZ, AA, WJC, S-PN, PCD, and MR wrote the paper. EPS, WJC, SG-T, MBL, RDP, PCD and MR provided supervision and funding support.

## Conflict of interest

PCD is on the scientific advisory board to Sirenas, Cybele, Microbiome, and Galileo.

## Notes

https://gnps.ucsd.edu/ProteoSAFe/status.jsp?task=525fd9b6a9f24455a589f2371b1d9540.

http://gnps.ucsd.edu/ProteoSAFe/status.jsp?task=e2bd16458ec34f3f9f99982dedc7d158.

https://gnps.ucsd.edu/ProteoSAFe/status.jsp?task=525fd9b6a9f24455a589f2371b1d9540

http://gnps.ucsd.edu/ProteoSAFe/status.jsp?task=e2bd16458ec34f3f9f99982dedc7d158

