## Supplementary Information for "Siderophore-mediated zinc acquisition enhances enterobacterial colonization of the inflamed gut"

**Supplemental information.**

**Allelic exchange deletion of *znuA* in *E. coli* Nissle 1917**

To construct an *E. coli* Nissle 1917 (EcN; **Supplementary** **Table 1**) *znuA* mutant, DNA regions of approximately 600-800 bp in length flanking the *znuA* gene were amplified by PCR with primers *znuA* FR1-Fw and *znuA* FR1-Rv (upstream flanking region, FR1) and primers *znuA* FR2-Fw and *znuA* FR2-Rv (downstream flanking region, FR2). A *Bam*HI restriction site was added to the 5’ end of primers *znuA* FR1-Fw and *znuA* FR2-Rv, and an *Xba*I restriction site was added to the 5’ end of primers *znuA* FR1-Rv and *znuA* FR2-Fw (**Supplementary** **Table 2**). The blunt-end flanking region PCR products were digested with XbaI, then ligated together using the Quick Ligation kit (New England Biolabs). Phusion High Fidelity DNA polymerase and primers *znuA* FR1-Fw and *znuA* FR2-Rv were used to amplify the FR1 + FR2 ligation product (FR1-FR2). A PCR product of the predicted size was gel-purified and ligated into pCRBlunt II-TOPO using the ZERO Blunt cloning kit (Invitrogen). The construct was heat-shocked into *E. coli* TOP10 and plated on LB+Kan plates. Plasmids isolated from single colonies were screened by *Eco*RI digestion, and positive clones were confirmed by sequencing using M13 Fw and M13 Rv universal primers. Accurate clones were designated pNM3 (pCRBlunt II-TOPO::*znuA* FR1-FR2). The plasmid pNM3 was digested with BamHI, and the FR1-FR2 fragment was gel-purified, ligated to BamHI-digested suicide vector pRDH10, then introduced into chemically competent *E. coli* CC118 λpir cells by heat-shock followed by selection on LB+Cm agar. A positive clone was designated pNM4 (pRDH10::*znuA* FR1-FR2). To enable conjugation of pNM4, purified plasmid was heat-shocked into *E. coli* S17-1 λpir. Wild-type EcN carrying the temperature-sensitive plasmid pSW172 (Lopez et al., 2012) was conjugated with *E. coli* S17-1 λpir containing pNM4 on LB agar at 30 °C (to allow for pSW172 replication). Single-crossover transconjugants were then selected for on LB+Carb+Cm agar. Afterwards, the transconjugants were subjected to sucrose selection in order to counterselect cells still harboring the integrated pNM4 plasmid, thus yielding WT revertants or Δ*znuA* mutants. Mutants were confirmed by PCR, then pSW172 was cured by growth at 37 °C. The resulting mutant strain was termed JZL95 (EcN Δ*znuA*(-82 to +1000)). An EcN Δ*znuA*::KSAC strain was also generated (JZL109), first cloning the *Xba*I-digested KSAC kanamycin resistance cassette from pBS34 into the *Xba*I restriction site of pNM4, yielding pNM4::KSAC (pRDH10::*znuA* FR1-KSAC-FR2).

**Construction of EcN *znuA zupT* mutant**

The EcN *znuA* *zupT* mutant was constructed using the lambda Red recombinase system (Datsenko and Wanner, 2000) . Briefly, primers (*zupT*-Fw and *zupT*-Rv) were designed with sequences homologous to the 5′ and 3′ ends of the EcN *zupT* gene and sequences homologous to the kanamycin resistance cassette of pKD4. The primers were used to PCR amplify the kanamycin resistance cassette. The PCR product was gel-purified, then electroporated into EcN *znuA* (JZL95) carrying pJK611 (a sucrose-counterselectable variant of pKD46). Kanamycin and sucrose were used for selection of deletion mutants and counterselection of pJK611, respectively. Putative mutants were screened on LB+Carb agar to confirm loss of pJK611, and the mutation was then confirmed using PCR. The resulting strain was termed JZL100 (EcN Δ*znuA*(-82 to +1000) Δ*zupT*(-42 to +774)::Kan).

**Construction of EcN *ybtX, znuA ybtX*, and *znuA zupT ybtX* mutants**

EcN *znuA* *ybtX* and EcN *znuA* *zupT* *ybtX* were constructed by allelic exchange, employing EcN *znuA* (JZL109) and EcN *znuA zupT* (JZL100) as parental strains. Using the NEBuilder tool (http://nebuilder.neb.com), overlapping primers were designed to construct a deletion vector for the *ybtX* gene (-108 to +1281). 1000 bp upstream and downstream of the EcN *ybtX* gene, as well as the chloramphenicol resistance cassette from pKD3, were amplified with the specific primers. Suicide plasmid pGP704 was digested with *Eco*RV and *Sal*I. PCR products and plasmids were gel-purified, then equimolar concentrations were used in a Gibson Assembly reaction, then transformed into *E. coli* CC118 λpir, selecting on LB+Cm agar. Clones were screened by sequencing the plasmid insert, and an accurate clone was designated pJB10 (pGP704::*ybtX* FR1-Cm-FR2). pJB10 was then electroporated into *E. coli* S17-1 λpir. EcN *znuA* and EcN *znuA zupT* were then separately conjugated with *E. coli* S17-1 λpir pJB10 on LB agar, after which transconjugants were selected for by plating on LB+Kan+Cm agar. Double-crossover mutants were identified by patching colonies on LB+Carb and LB+Cm plates. Carb^S^ Cm^R^ colonies were checked by PCR to assess for the loss of *ybtX* (ybtX_pres_Fw and ybtX_pres_Rv) and the presence of the chloramphenicol resistance cassette (Gib_out_LB_Fw and C2; Gib_out_RB_Rv and C1). The resulting strain EcN *znuA zupT ybtX* was termed JB76, while EcN *znuA ybtX* was termed JB92.

The EcN *ybtX* mutant was constructed using the lambda Red system. Briefly, primers (ybtX_red_Fw and ybtX_red_Rv2) were designed with sequences homologous to the 5′ and 3′ ends of the EcN *ybtX* gene and to the chloramphenicol resistance cassette of pKD3. The primers were used to amplify the chloramphenicol resistance cassette by PCR. The PCR product was electroporated into EcN wild-type carrying pJK611. Chloramphenicol and sucrose were used for selection of deletion mutants and counterselection of pJK611, respectively. The mutation was confirmed using PCR, and pJK611 was confirmed to be cured by plating on LB+Carb agar. The resulting strain was termed JB90 (EcN Δ*ybtX*(-108 to +1281)::Cm).

**Construction of EcN *irp2* and EcN *znuA zupT irp2* mutants**

A mutant in EcN carrying a deletion of the *irp2* open reading frame was constructed using the lambda Red system. Briefly, primers (irp2up-TetRA F and irp2dn-TetRA R) were designed with sequences homologous to the 5′ and 3′ ends of the EcN *irp2* gene and to the *tetRA* resistance cassette of MSC74. The primers were used to amplify the tetracycline resistance cassette by PCR. The PCR product was electroporated into wild-type EcN containing pJK611. Tetracycline and sucrose were used for selection of mutants and counterselection of pJK611, respectively. The *irp2* deletion was confirmed using PCR (irp2inF and irp2inR) and sequencing (irp2-upF and irp2-dnR), and also confirmed for the loss of plasmid pJK611 by plating on LB+Carb plates. The resulting strain was termed HZE116 (EcN Δ*irp2*::Tet).

A deletion of the *irp2* open reading frame in EcN *znuA* *zupT* (JZL100) was constructed by allelic exchange. Using the NEBuilder tool (http://nebuilder.neb.com), overlapping primers were designed to construct a deletion vector for the *irp2* gene. 500bp upstream (pGP704-salI US irp2 F and TetR-US irp2 R) and downstream (TetA-DS irp2 F and pGP704-sacI DS-irp2 R) regions of EcN flanking the *irp2* gene, as well as the tetracycline resistance cassette (TetR-F and TetA-R) of MSC74, were then amplified. Suicide plasmid pGP704 was digested with SalI and SacI. PCR products and plasmid were gel-purified, then equimolar concentrations were used in a Gibson Assembly reaction, followed by transformation into DH5α λ*pir* and selection on LB+Tet agar. Clones were screened by sequencing, and an accurate clone was designated pHZE107 (pGP704::*ybtX* FR1-Tet-FR2). pHZE107 was then purified and electroporated into *E. coli* S17-1 λpir. EcN *znuA zupT* was conjugated with *E. coli* S17-1 λpir pHZE107 on LB agar. Transconjugants were then selected for by plating on LB+Kan+Tet, and double-crossovers were identified by screening for Carb^S^ colonies. Kan^R^ Tet^R^ Carb^S^ colonies were then tested by PCR with primers checking for the loss of *irp2* (irp2inF and irp2inR) and the presence of the tetracycline resistance cassette (TetR-F and TetA-R). The resulting strain was termed HZE112 (EcN *znuA zupT irp2*).

**TABLES**

**Supplementary Table 1**

| **Strain or plasmid** | **Relevant characteristics** | **Source or Reference** |
| --- | --- | --- |
| *Escherichia coli* strains | | |
| CC118 λ*pir* | F- *araD139* Δ(*ara, leu*)7697 ∆*lacX74* *phoA*∆20 *galE* *galK* *thi* *rpsE rpoB argE*_am_ *recA1* λ*pir* | (Herrero et al., 1990) |
| DH5α λ*pir* | F- *supE44* Δ*lacU169* φ80d*lacZ*ΔM15 *recA1* *endA1* *hsdR17* *thi-1* *gyrA96* *relA1* λ*pir* | Lab strain |
| DH5α MCR | F- *mcrA* Δ(*mrr-hsdRMS-mcrBC*) φ80d*lacZ*ΔM15 Δ(*lacZYA-argF)*U169 *deoR* *recA1 endA1 phoA supE44* λ- *thi-1 gyrA96 relA1* | (Grant et al., 1990) |
| S17-1 λ*pir* | F- *recA* *thi* *pro* r_K_^-^ m_K_^+^ RP4::2-Tc::*Mu*Km Tn7 λ*pir* | (Herrero et al., 1990) |
| TOP10 | F- *mcrA* Δ(*mrr-hsdRMS-mcrBC*) φ80d*lacZ*ΔM15 Δ*lacX74* *recA1 araD139* Δ(*ara-leu*)7697 *galU galK* *rpsL*(Str^R^) *endA1 nupG* λ- | Invitrogen |
| EcN | *E. coli* Nissle 1917 wild-type | ArdeyPharm, Germany |
| JZL95 (*znuA*) | EcN Δ*znuA*(-82 to +1000) | This study |
| JZL109 (*znuA*) | EcN Δ*znuA*(-82 to +1000)::Kan | This study |
| JZL100 (*znuA zupT*) | EcN Δ*znuA*(-82 to +1000) Δ*zupT*(-42 to +774)::Kan | This study |
| JB90 (*ybtX*) | EcN Δ*ybtX*(-108 to +1281)::Cm | This study |
| JB92 (*znuA ybtX*) | EcN Δ*znuA*(-82 to +1000)::Kan Δ*ybtX*(-108 to +1281)::Cm | This study |
| JB76 (*znuA zupT ybtX*) | EcN Δ*znuA*(-82 to +1000) Δ*zupT*(-42 to +774)::Kan Δ*ybtX*(-108 to +1281)::Cm | This study |
| HZE116 (*irp2*) | EcN Δ*irp2*(+1 to +6108*)*::Tet | This study |
| HZE112 (*znuA zupT irp2*) | EcN Δ*znuA*(-82 to +1000) Δ*zupT*(-42 to +774)::Kan Δ*irp2*(+1 to +6108*)*::Tet | This study |
| *Salmonella enterica* serovar Typhimurium strains | | |
| IR715 | ATCC 14028 Nal^R^ | (Stojiljkovic et al., 1995) |
| JZL3 | IR715 Δ*znuA*::Cm | (Liu et al., 2012) |
| APJ4 | IR715 Δ*znuA*::Cm Δ*zupT*::KSAC | (Cerasi et al., 2013) |
| MSC74 | IR715 Δ*iroN*::*tetRA* | (Sassone-Corsi et al., 2016) |
| Plasmids | | |
| pBS34 | pBluescriptII KS+ ::[XbaI][PstI]KSAC[PstI][XbaI],  Carb^R^ Kan^R^ | (Raffatellu et al., 2009) |
| pCRBlunt II-TOPO | TA-Cloning Vector, Kan^R^ | Invitrogen |
| pGP704 | *oriR6K mobRP4*, Carb^R^ | (Miller and Mekalanos, 1988) |
| pHP45Ω | Strep^R^ Carb^R^ | (Prentki and Krisch, 1984) |
| pKD3 | Carb^R^ Cm^R^ | (Datsenko and Wanner, 2000) |
| pKD4 | Carb^R^ Kan^R^ | (Datsenko and Wanner, 2000) |
| pJK611 | pKD46::*sacB* | Kelly T. Hughes |
| pJB10 | pGP704::*ybtX* FR1-Cm-FR2, Cm^R^, Carb^R^ | This study |
| pNM3 | pCRBlunt II-TOPO::*znuA* FR1+FR2, Kan^R^ | This study |
| pNM4 | pRDH10::*znuA* FR1-FR2, Cm^R^ | This study |
| pNM4::KSAC | pRDH10::*znuA* FR1-KSAC-FR2, Cm^R^ Kan^R^ | This study |
| pRDH10 | *oriR6K mobRP4 sacRB*, Cm^R^ Tet^R^ | (Kingsley et al., 1999) |
| pSW172 | *oriR101* *repA101ts*, Carb^R^ | (Lopez et al., 2012) |
| pHZE107 | pGP704::*irp2* FR1-Tet-FR2, Tet^R^ Carb^R^ | This study |

**Supplementary Table 2**

| **Name** | **Sequence (5' - 3')** | **Source** |
| --- | --- | --- |
| *znuA* FR1-Fw | CACATTGGATCCGTTGTTCAGCACCACGAG | This study |
| *znuA* FR1-Rv | TATGCCTCTAGACAAGTCTGGTTTCCCTGG | This study |
| *znuA* FR2-Fw | TATGCCTCTAGACACACCGCGTTATGTTGG | This study |
| *znuA* FR2-Rv | CACATTGGATCCCGGAACTGTTTCGCCGTC | This study |
| *zupT*-Fw | GTAAGAACCCGGATAACAATGATGATGATCATCAGTTATTGTGTAGGCTGGAGCTGCTTC | This study |
| *zupT*-Rv | TGTTGCCTTTAGCAATGGGCAACATCTGTCATTATCGTCTATGGGAATTAGCCATGGTCC | This study |
| LB_ybtX_fwd | gccacctgcagatctgcaggCAGACCCCAGATGCTGAAC | This study |
| LB_ybtX_rev | gtcacaggtaTCCGCGAATAACACAGAG | This study |
| CmR_fwd | tattcgcggaTACCTGTGACGGAAGATCACTTCGC | This study |
| CmR_rev | cagggtactcCTTACGCCCCGCCCTGCC | This study |
| RB_ybtX_fwd | ggggcgtaagGAGTACCCTGCTCAACCACCTGTCCCTC | This study |
| RB_ybtX_rev | aattcccgggagagctcgatAGCCGGGCGCTGCTGGCG | This study |
| ybtX_pres_FW | AAAGAGGGTGAAGTCGACAC | This study |
| ybtX_pres_RV | GTCTATGCAGTCCTTACCCG | This study |
| ybtX_LB_FW | CTGACGGAACATAAACGAGC | This study |
| ybtX_RB_RV | TACAGGTGGTGGTGTTATCC | This study |
| Gib_out_LB_FW | CAATTTGCATAATCGCGTTCAG | This study |
| Gib-out-RB-RV | AGGGGATGATGGTATTGCGC | This study |
| ybtX_RED_FW | CACTGATTTTCATCCCTTACCTCTCTGTGTTATTCGCGGAGTGTAGGCTGGAGCTGCTTC | This study |
| ybtX_LB_FW | CTGGTGATGGAAGAGGGACAGGTGGTTGAGCAGGGTACTCCATATGAATATCCTCCTTA | This study |
| Gib-out-RB-RV | AGGGGATGATGGTATTGCGC | This study |
| ybtX_RED_FW | CACTGATTTTCATCCCTTACCTCTCTGTGTTATTCGCGGAGTGTAGGCTGGAGCTGCTTC | This study |
| ybtX_LB_FW | CTGGTGATGGAAGAGGGACAGGTGGTTGAGCAGGGTACTCCATATGAATATCCTCCTTA | This study |
| TetR-F | TTAAGACCCACTTTCACATTTAAG | This study |
| TetA-R | CTAAGCACTTGTCTCCTGTTTAC | This study |
| pGP704-salI US irp2 F | acctgcagatctgcaggtcgacTCAGTCTGGTGCTGGATG | This study |
| TetR-US irp2 R | AATGTGAAAGTGGGTCTTAATCTTCCTCCTGATGGCACG | This study |
| TetA-DS irp2 F | AACAGGAGACAAGTGCTTAGCGCGAAGCAAACTGATTTTCC | This study |
| pGP704-sacI DS-irp2 R | tcgaattcccgggagagctcTGGCAATATAGTCTTTATCATTG | This study |
| irp2up-TetRA F | ATGCTTTTCGGTAAGACGTGCCATCAGGAGGAAGATTAAGACCCACTTTCACATTTAAG | This study |
| irp2dn-TetRA R | GATGGCGTTCCGGGGAAAATCAGTTTGCTTCGCGCTAAGCACTTGTCTCCTGTTTAC | This study |
| irp2-inF | ACAACGCTTCCTCGTACAATG | This study |
| Irp2-inR | TCTGTTCGAGCACCTGTTGC | This study |
| Irp2-upF | TTTGGCGTAACTCCTTCGAC | This study |
| Irp2-dnR | TTCATAAAGTCAATGGAACG | This study |

**Supplementary figures**

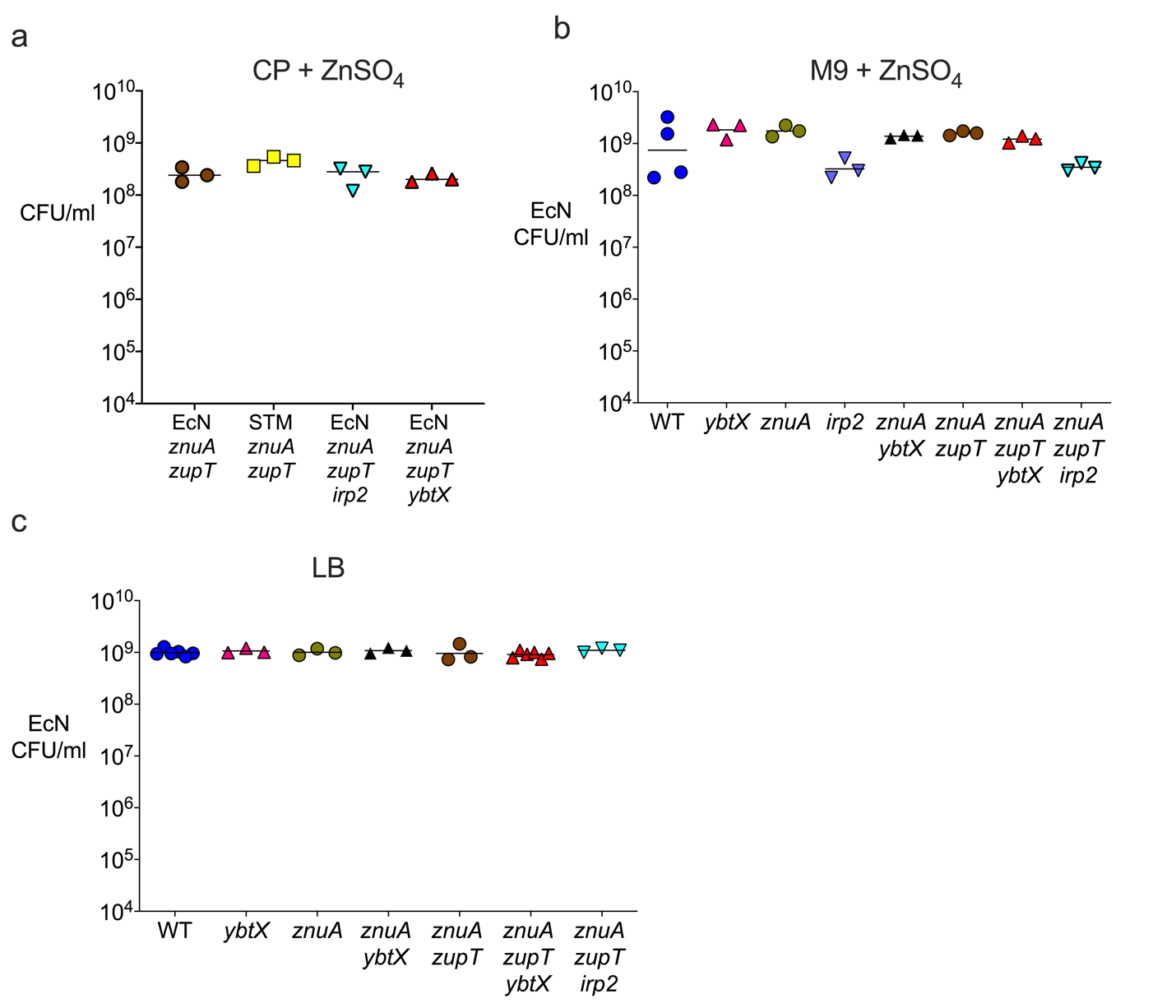

**Supplementary Figure 1.** **Wild-type and mutant strains grow to similar abundance in rich media or in zinc-limiting media supplemented with zinc.**

(**a**) EcN and STm strains were grown in modified LB medium containing 150 µg/ml CP and 5 µM ZnSO_4_ for 16 h static incubation at 37 °C. (**b**) EcN strains were grown in M9 medium supplemented with 5 µM ZnSO_4_ for 20 h shaking incubation at 37 °C. (**c**) EcN strains were grown in LB medium for 20 h shaking incubation at 37 °C. Each experiment was done independently three times.

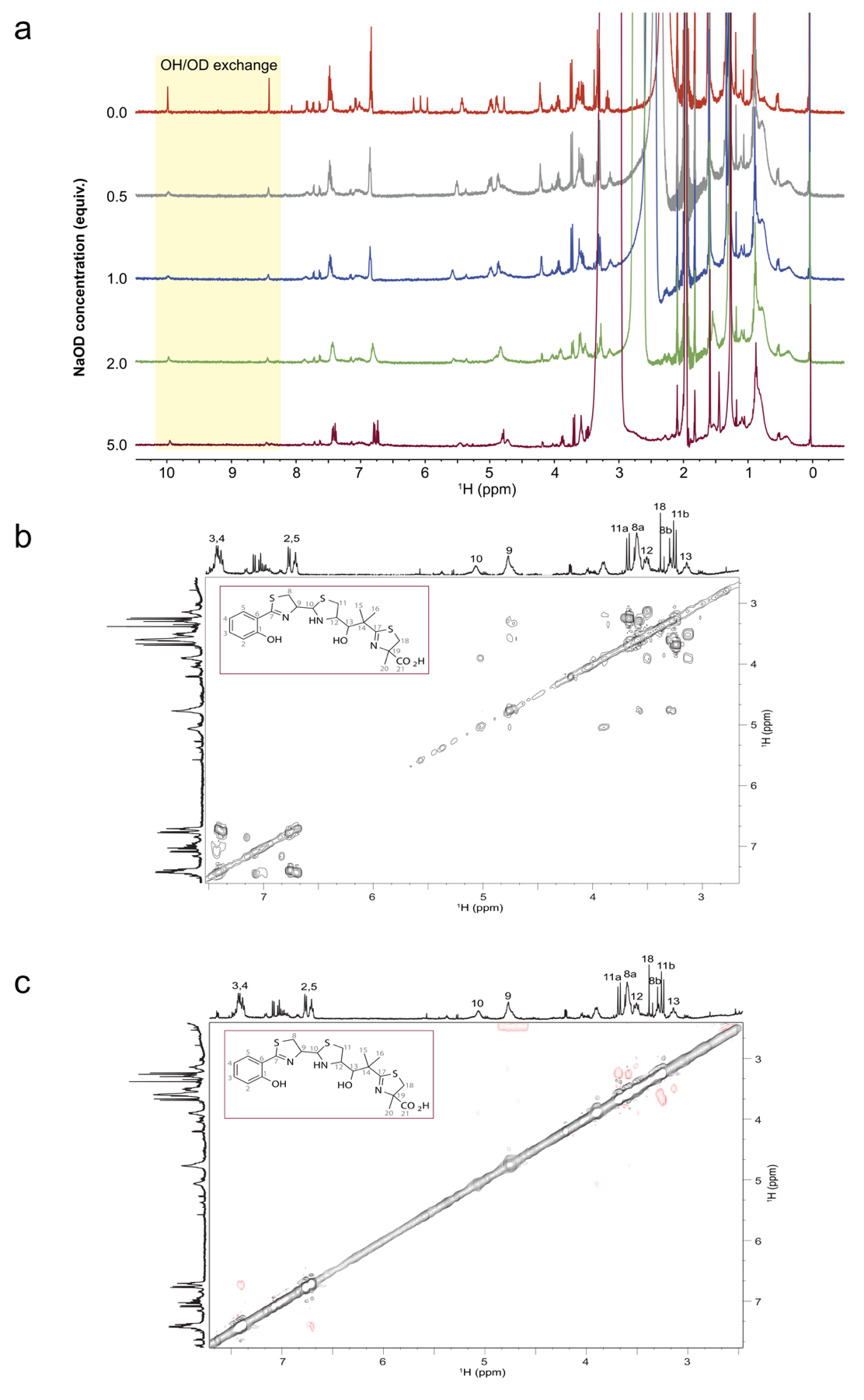

**Supplementary Figure 2. Verification of NMR resonance assignments of Ybt by 2D NMR, and lack of significant changes in Ybt resonances upon the addition of base**.

(**a**) ^1^H NMR 1D spectra of Ybt dissolved in CD_3_CN in the presence of 5.0 equiv. of ZnCl_2_ (top trace, 0 equiv. of NaOD, red trace) and increasing amounts of NaOD (0.5 equiv., gray trace), (1.0 equiv., blue trace), (2.0 equiv., green trace), and (5.0 equiv., brown trace). There are not changes in the spectrum other than the loss of OH signal intensity upon addition of NaOD (highlighted in yellow), which is attributed to increased solvent exchange. The starting spectrum of Ybt in the presence of 5.0 equiv. ZnCl_2_ shows hydrolysis products (signal at 6.2 ppm) in addition to bound and unbound Ybt isomers. (**b**) 2D gradient-enhanced ^1^H-^1^H COSY showing aliphatic and aromatic region of spectrum for Ybt dissolved in CD_3_CN. (**c**) 2D gradient-enhanced ^1^H-^1^H ROESY showing aliphatic and aromatic region of spectrum for Ybt dissolved in CD_3_CN. All spectra were acquired at 500 MHz.

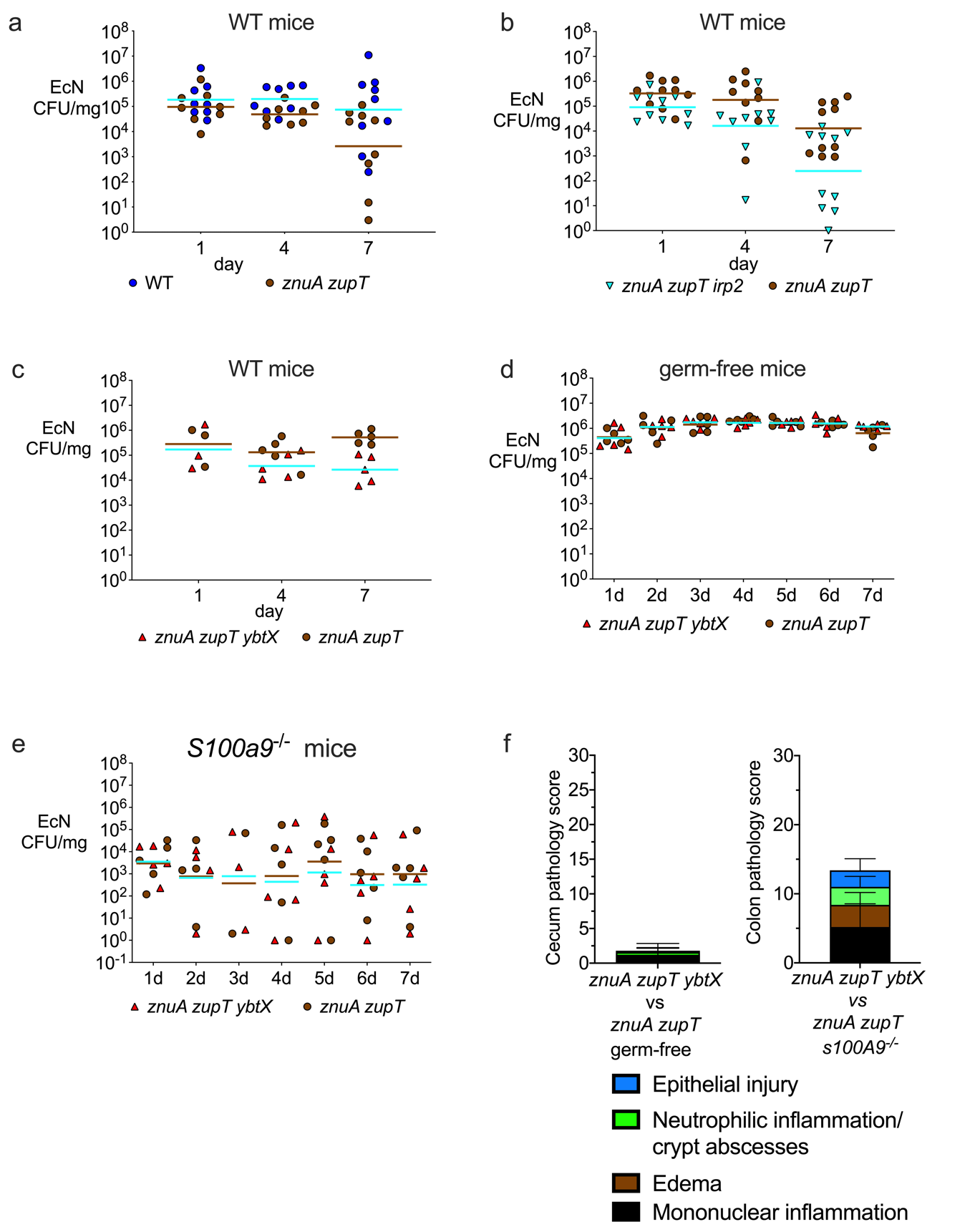

**Supplementary Figure 3.** **EcN fecal CFU and pathology scores from mice in Figures 4 and 5.**

(**a-e**) CFU of EcN strains in the fecal content of (**a-c**) wild-type C57BL/6 mice treated with DSS, (**d**) wild-type germ-free Swiss Webster mice, (**e**) C57BL/6 *S100a9^-/-^* mice, all inoculated with the indicated 1:1 mixtures of EcN strains. Brown bars represent the geometric mean of the *znuA zupT* mutant in each experiment, blue bars represent the geometric mean of the other strain represented in the figure panel. (**f**) Pathology scores of cecal tissue from day 7 germ-free mice in panel D (left chart), or of colonic tissue from day 7 *S100a9^-/-^* DSS-treated mice in panel E (right chart).
